# De-novo design of a random protein walker

**DOI:** 10.1101/2025.09.29.677966

**Authors:** Liza Ulčakar, Hao Shen, Eva Rajh, Tadej Satler, Federico Olivieri, Joseph Watson, Yang Hsia, Justin Decarreau, Eric Lynch, Justin Kollman, David Baker, Ajasja Ljubetič

## Abstract

Molecular machines hold great potential. Design of static monomeric and oligomeric protein structures has advanced tremendously, but few dynamic protein systems have been designed. Here we present the design and characterization of a random protein walker that diffuses along a designed protein track. For the track, we designed micro-meter long fibres and developed a method to rigidly decorate them with arbitrary proteins. The walkers consist of homo-oligomers that reversibly bind the track using heterodimeric feet; we tested multiple heterodimer interfaces for reversibility and designed six walkers with different numbers of feet. Cryo-EM experiments confirmed the structure of the track and walkers. We performed detailed single molecule tracking and kinetics characterisation and found that the walkers with more feet diffuse along the track faster. The system represents a tuneable starting point for future powered protein molecular robots.

## Introduction

Molecular walkers, which convert energy into unidirectional motion, lie at the fascinating crossroads of biology, chemistry, and physics. Natural molecular motors such as dynein^1^, kinesin^2^, and myosin^3^ are essential for life and have been studied in detail^4^, but our understanding of them is still not complete. Programmable and robust *de novo* designed protein nanorobots could revolutionize fields from precision medicine to material science. Despite remarkable recent advances in *de novo* protein design^5^, building such functional systems remains out of reach. The state of the art includes a protein walker assembled from natural DNA binding proteins that can make steps along DNA with the help of microfluidics^6^ and *de novo* designed rotor proteins^7^. Tracking the motion of the rotors proved challenging, limiting their utility and further engineerability. Design of a random protein walker that moves along large distances would mark a substantial advance, deepening our understanding of molecular machine design and laying the groundwork for future protein-based nanodevices.

We set out to design a completely *de novo* designed protein system consisting of walkers with four to eight feet that specifically bind to a designed protein track with footholds (Fig. 1a). For conceptual simplicity, these walkers do not consume fuel; instead, their multivalent architecture allows individual feet to dynamically dissociate from and bind to successive footholds on the track. Our primary objective was to enable and characterize the diffusion of the walker along the track, thereby probing how multivalency influences both speed and processivity. This minimal system provides a versatile scaffold onto which external energy sources could be integrated in future designs.

**Fig. 1:**
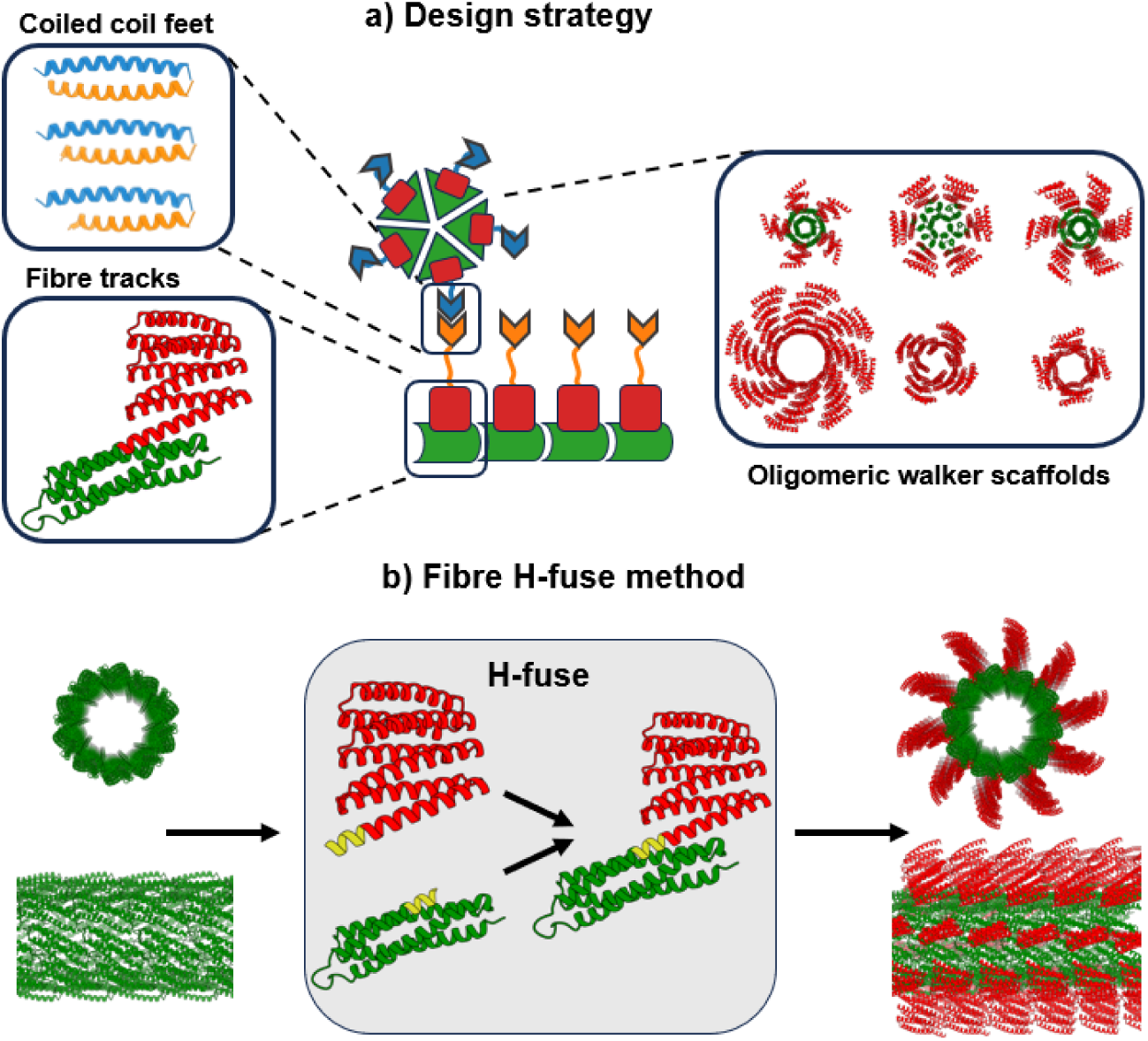
Essential parts for a random walker system. **(a)** Schematics for the random walker system. The tracks are self-assembled from protomers, consisting of a four helical bundle (green), a rigidly attached DHR (red) and a flexibly attached P3SN_4h, P3SN_3.5h and P3SN_3h (orange). The walkers are cyclic oligomers with a helical bundle core (green), rigidly fused DHRs (red) and flexibly fused P4SN (blue). **(b)** Strategy for attaching a DHR to the fibre using the H-fuse method. A structural overlap between the C-terminus of the four helical bundle and the N-terminus of a DHR is identified and the structures are rigidly fused into a single-chain. The residues in the junction are redesigned to prevent steric clashes.

## Results

We reasoned that, at a minimum, three components would be required to construct a random walker: (i) long protein fibres to serve as tracks; (ii) symmetric protein scaffolds to act as the walkers; and (iii) well-behaved reversible heterodimers to function as feet and footholds, forming the critical interfaces that enable stepping along the track.

### Design strategy and structural characterization of the tracks

Protein fibres that self-assemble from globular monomers have been successfully designed^8^, however they were too short to be distinctly resolved as fibres with total internal reflection fluorescence (TIRF) microscopy, a crucial technique for single molecule tracking experiments. To address this limitation, we aimed to design fibres with a larger diameter to increase their rigidity, with the goal of further promoting fibre length and structural stability.

The core of the tracks was designed by docking protein scaffolds and redesigning the interface^8^; 17 scaffolds were used. We designed 9437 symmetry configurations sampling a wider range of cyclic and N jumps (allowing the basic helical transform to repeat N times before contact occurs) combinations for larger diameter. 74 designs were tested experimentally and two constructs, HB5 and HB67, formed long fibres. The HB67 fibre was expressed in fusion with GFP and imaged with a confocal microscope (Extended Data Fig. 1). The length of the fibres was in the micrometre range. However, their structure could not be resolved by cryo-electron microscopy (cryo-EM) due to heterogeneous fibre diameter.

We reasoned that additional structural markers could facilitate structure determination. The previously described H-fuse method^9^ can rigidly fuse two protein domains using overlapping secondary structure, however the method is not compatible with fibres. We therefore developed **Fibre H-fuse,** a new approach for rigidly attaching arbitrary proteins to fibrous assemblies (Fig. 1b). Using *in silico* design, we generated fusions of the fibre protomer with 53 distinct designed helical repeat proteins (DHRs)^10^, scanning several overlap positions per DHR. Constructs that showed favourable secondary-structure overlap and lacked internal or symmetric clashes were further refined by redesigning the fusion junction. From this set, 18 final models were selected based on Rosetta scoring metrics and manual inspection, and corresponding synthetic genes were ordered for experimental verification. Two of the proteins – HA4 and HA13 – both expressed in a soluble form and formed fibres, which was confirmed by confocal microscopy (Extended Data Fig. 2). HA13 was further structurally characterized by cryo-EM to 3.3 Å resolution (Fig. 2a, Extended Data Fig. 3). The structure revealed a fibre with an inner diameter of 10 nm and the longitudinal spacing of 4.7 nm between the attached DHRs. The experimentally determined structure has a smaller radius than the design (Extended Data Fig. 3e).

**Fig. 2:**
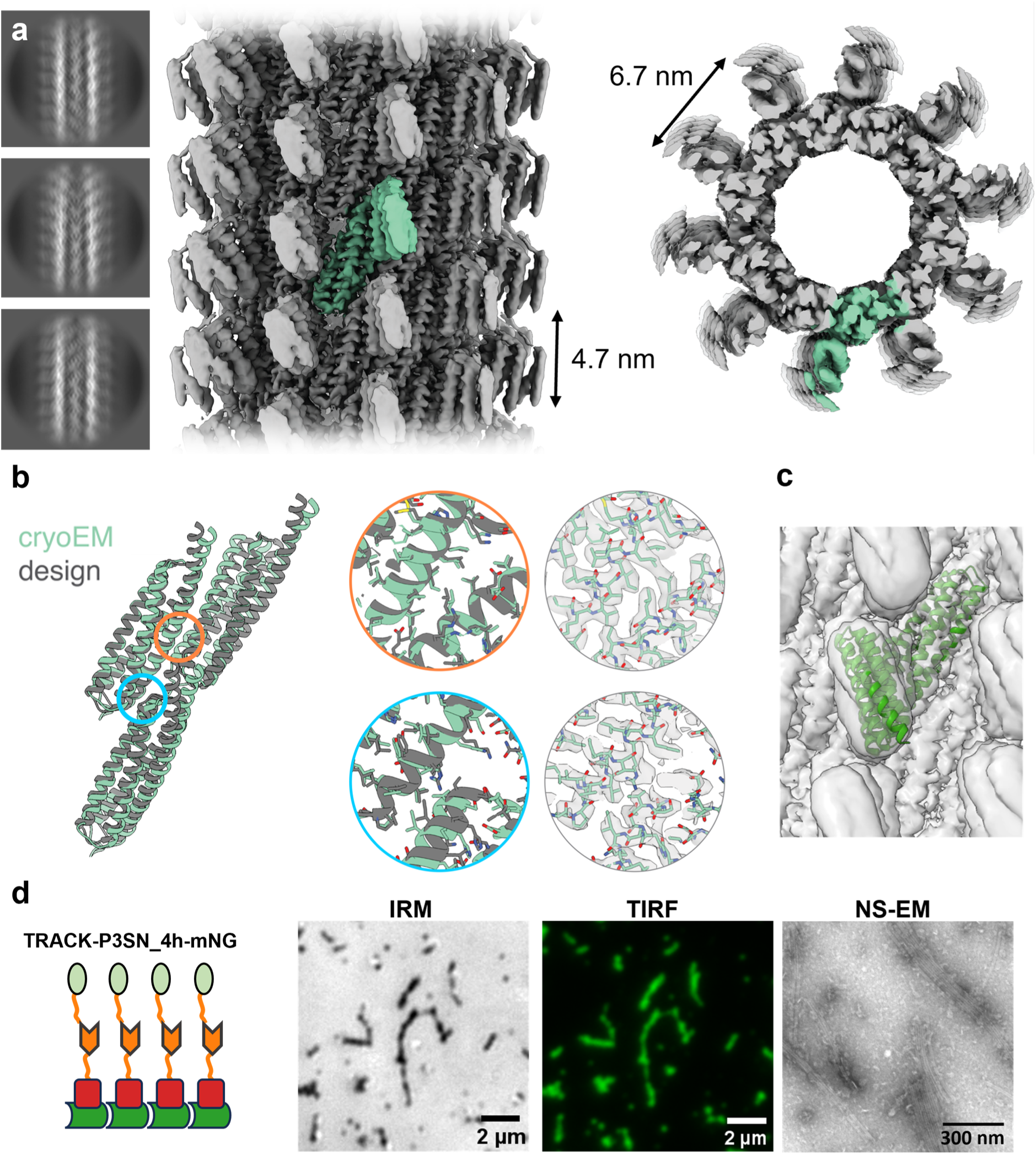
Structural characterization of the tracks. **(a)** Cryo-EM reconstruction of the fibre HA13. Left, representative 2D class averages. Middle, side view of the reconstructed density map showing the axial spacing (4.7 nm) between adjacent footholds; a single protomer is highlighted in green. Right, top view revealing the helical arrangement and the perpendicular spacing (6.7 nm) between protomers. **(b)** Overlay of design (grey) and cryo-EM model (green) highlighting the close match between the designed and experimentally determined structures. Insets show close-up views of inter-protomer interfaces (orange and blue circles) alongside the corresponding fit into the density map. **(c)** Cryo-EM density of the fibre (grey) overlaid with a design cartoon model (green). **(d)** Experimental validation of track assembly. Left, schematic of the fibre design (fibre, dark green; DHR, red; footholds, orange; mNG, light green), followed by IRM, TIRF and NS-EM images. Micrographs for all tracks are shown in Extended Data Fig. 4.

In addition to facilitating electron microscopy, the fusion of DHRs also provide additional spacing from the fibre interface, reducing the chance of perturbing the fibre’s assembly upon adding the footholds for the walker.

### Selection of heterodimeric feet/footholds

To complete the track, we required heterodimer protomers capable of binding and unbinding to serve as footholds on the fibre. Initial testing with a shorter heterodimeric four helix bundle mALb8^11^ showed that presence of footholds can interfere with fibre assembly. We evaluated 16 rigid and 4 flexible fusions of mALb8 to the fibre, but none of these constructs successfully formed fibres. We hypothesised that this failure was due to off-target homodimerization of the foothold domains. To address this, we adopted a high-throughput screening strategy, creating a system to flexibly attach any heterodimeric protomer to the fibre. Footholds for the walker were linked to the fibre using a flexible 10 amino acid linker. A total of 46 constructs (23 heterodimers on both HA4 and HA13) were screened, including heterodimeric four-helical bundles^12^, designed parallel and antiparallel coiled-coils^13^ (CCs), as well as large designed heterodimers^14^ (LHDs, Extended Data Table 1).

Among the tested heterodimers, parallel CCs and LHDs demonstrated the most favourable properties in terms of fibre assembly, leading us to select the P3:P4 coiled-coil pair^13^. The soluble variant of P3 (P3SN) was used as the foothold on the fibre, while P4SN served as the foot of the walkers. To investigate how walker–track affinity influences walker movement, we designed three track variants that differed in the length of the P3SN coiled-coil domain, containing either 4, 3.5, or 3 heptad repeats; shorter coils were expected to exhibit weaker binding. For an overview of designed tracks and walkers see Extended Data Table 2.

To facilitate imaging, the track subunits were fused to the fluorescent protein mNeonGreen (mNG) via a 50-amino-acid flexible linker. However, due to concerns that the attached mNG might sterically hinder walker movement, we also characterised track variants lacking the fluorescent fusion. All tracks were visualised using total internal reflection fluorescence (TIRF) microscopy, interference reflection microscopy (IRM), and negative stain electron microscopy (NS-EM). An example track is shown in Fig. 2d; all others are shown in Extended Data Fig. 4.

### Walker design

Next, we sought protein scaffolds capable of accommodating variable numbers of feet. We selected *de novo*–designed symmetric oligomers with cyclic symmetries ranging from C4 to C8^15,16^ (Fig. 1a, Extended Data Table 2, Extended Data Table 3). These oligomers were either assembled directly from DHRs or had DHRs fused to their cores. To ensure that the spacing between termini matched the fibre’s axial repeat of 4.7 nm, we fine-tuned the length of the DHR segments by adding or removing repeats (Extended Data Table 3). Finally, we linked the P4SN coiled-coil foot to the scaffold via a flexible six–residue linker; P4SN specifically binds to the complementary P3SN coiled-coil footholds displayed on the fibre.

All six walker constructs were successfully expressed in *E. coli* and purified from the soluble fraction. The oligomeric assembly states were validated using both size-exclusion chromatography coupled to multi-angle light scattering (SEC-MALS) and mass photometry (MP), performed at micromolar and nanomolar concentrations, respectively (Fig. 3). Both techniques confirmed the expected molecular weights and oligomerization states of the designed assemblies. The MP histogram revealed potential additional species. Notably, a peak near the lower detection limit (30–40 kDa) was observed that most likely represent monomeric species or noise and impurities. Peaks corresponding to C5- and C7-sized assemblies were observed for WALKER-C6 and WALKER-C8, respectively, suggesting low levels of alternative oligomeric states.

**Fig. 3:**
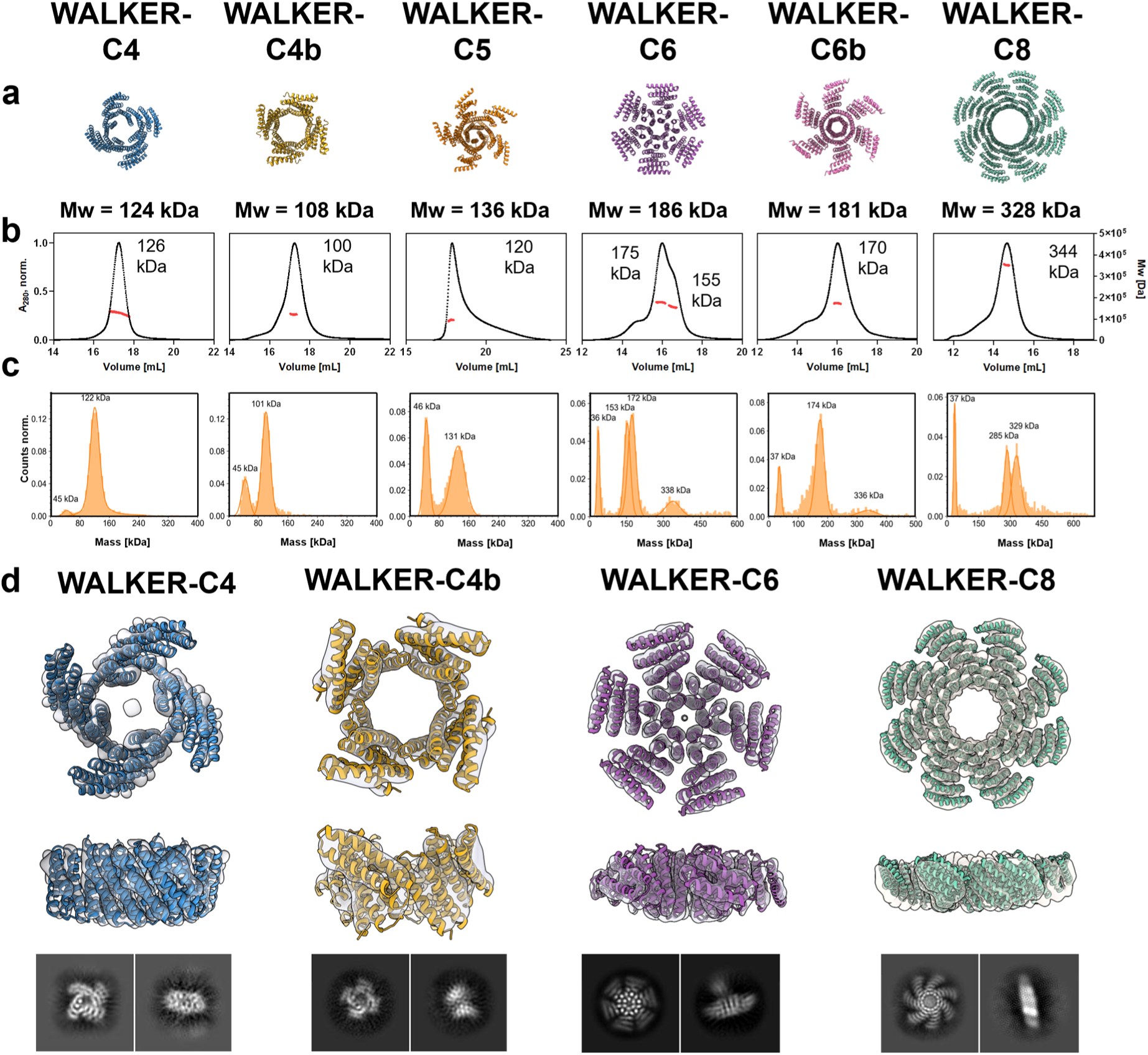
Biophysical characterization and structural validation of designed walker oligomers. **(a)** Computational models of the designed walker oligomers, labelled according to their symmetry (e.g., WALKER-C4, -C5, -C6, -C8), with corresponding theoretical molecular weights (Mw) indicated. **(b)** SEC-MALS reveals mostlymonodisperse species with experimentally determined molecular weights consistent with design expectations. (c) MP histograms confirming the oligomeric states of the walkers. **(d)** Cryo-EM density maps (grey surface) overlaid with atomic models (coloured cartoons) of selected walkers (C4, C4b, C6, and C8), shown in both top and side views. Representative 2D class averages are displayed below each structure.

To assess the stability of the walker assemblies under the low nanomolar concentrations required for single-molecule tracking, MP was performed two hours after dilution to low nanomolar concentrations of protomer (20 – 40 nM, depending on the walker) (Extended Data Fig. 5). All walkers except C4b retained their oligomeric state for at least two hours, which is longer than the duration of the typical single molecule tracking experiment. WALKER-C4b fully dissociated after two hours and was therefore excluded from subsequent single-molecule tracking analyses experiments. The walkers retain their oligomeric states after two hours at 95 °C (Extended Data Fig. 5).

The structures of walkers C4, C4b, C6 and C8 were determined by cryo-EM (Fig. 3d), achieving resolutions of 4.46 Å, 4.45 Å, 3.89 Å, and 4.12 Å, respectively. Due to the flexible linkers, the feet could not be resolved. The cryo-EM reconstructions confirmed the oligomeric states previously established by SEC-MALS and MP, and the resulting density maps closely matched the corresponding computational design models.

### Single molecule tracking of walkers on tracks

To investigate walker dynamics along the tracks, we employed TIRF microscopy and IRM (Fig. 4a). Tracks were visualized in the green channel using TIRF, and in parallel with IRM, which enables label-free imaging. Walkers were fluorescently labelled with ATTO 643 and imaged in the far-red channel via TIRF at a frame rate of 4 Hz (Fig. 4a, left).

**Fig. 4:**
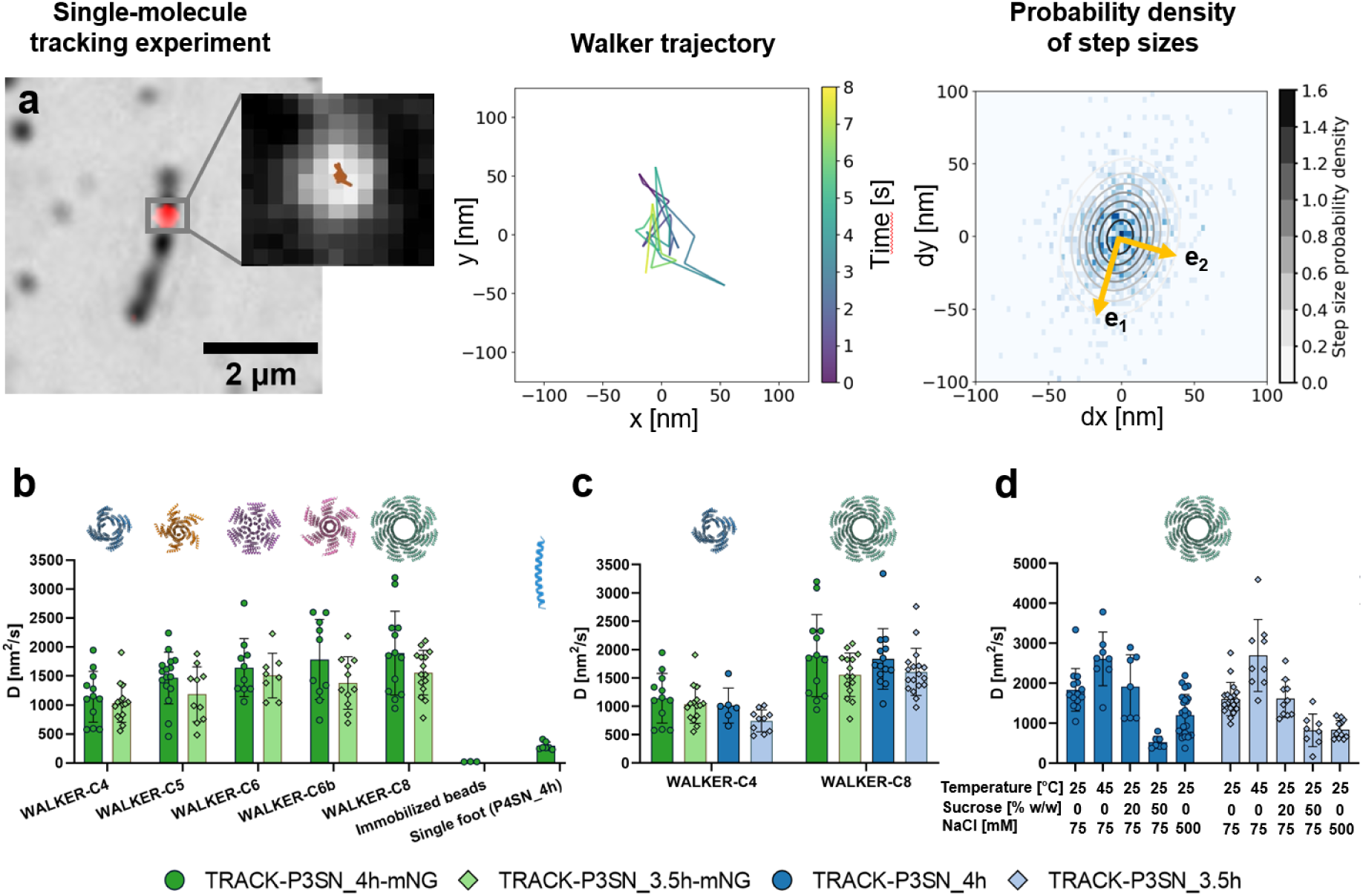
Single-molecule tracking and diffusion analysis of engineered random protein walkers. **(a)** Left: Representative TIRF micrograph showing a fluorescently labelled WALKER-C8 (red) moving along TRACK-P3SN_3.5h visualized via IRM. The inset shows the walker’s trajectory (brown) calculated from the single molecule data. Middle: A representative trajectory of a single WALKER-C8 on TRACK-P3SN_3.5h, with colours (purple to yellow) indicating time from the beginning of the particle’s trajectory. Right: Two-dimensional histogram of steps (dx, dy) pooled from all tracked walkers of a single recording. A fitted 2D Gaussian distribution reveals principal axes of motion (e_1_ and e_2_). Effective diffusion coefficients are calculated from the largest component, since it corresponds to motion along the track. **(b)** Diffusion coefficients of all tested walker variants (WALKER-C4 to WALKER-C8) measured on mNeonGreen labelled tracks with two different lengths of footholds (TRACK-P3SN_4h-mNG, dark green and TRACK-P3SN_3.5h-mNG, light green). Diffusion constants of immobilized beads and single feet (P4SN_4h) on TRACK-P3SN_4h-mNG served as negative controls. We observe that walkers with more feet diffuse faster. **(c)** Comparison of diffusion coefficients for two representative walker constructs (WALKER-C4 and WALKER-C8) on tracks with (TRACK-P3SN_4h-mNG, TRACK-P3SN_3.5h-mNG, green) and without (TRACK-P3SN_4h, TRACK-P3SN_3.5h, blue) mNeonGreen fusion. We observe that the mNeonGreen labelling does not impede the diffusion. **(d)** Diffusion coefficients of WALKER_C8 measured on unlabelled tracks (TRACK-P3SN_4h and TRACK-P3SN_3.5h) under varying environmental conditions: standard (25 °C, 0% sucrose, 75 mM NaCl), elevated temperature (45 °C, 0% sucrose, 75 mM NaCl), increased viscosity (25 °C, 75 mM NaCl, 20% and 50% sucrose) and increased salt concentration (25 °C, 0% sucrose, 500 mM NaCl). Higher temperatures, lower viscosity and lower salt concentration result in faster motion. Data in (b-d) represent mean ± s.d. from individual TIRF recordings.

Walker positions were localized by fitting the centre of the fluorescence intensity profiles using Picasso^17^ (Fig. 4a, inset). A concentration of 0.25 nM of walker protomer was used to ensure that each fluorescent spot corresponded to a single walker assembly. Individual positions were subsequently linked into trajectories using the Trackpy^18^ software package.

An example trajectory is shown in Fig 4a, middle, with further examples shown in Extended Data Fig. 6. Walkers are designed to move along the track axis, where the inter-foothold spacing is optimal (4.7 nm). Lateral steps perpendicular to the axis are also possible, albeit with less efficiency, due to the larger (6.3 nm) inter-foothold spacing. The anisotropy of the track is reflected in the trajectories: the walkers cover on average twice the distance along the fibre than perpendicular to the track (Extended Data Fig. 8, Extended Data Table 7). We measure average steps sizes of around 30 nm (Extended Data Table 5), suggesting that the individual steps could be faster than our camera framerate.

From the walker trajectories, we extracted a distribution of step sizes, which was fitted to a two-dimensional Gaussian function (Fig. 4a, right, Extended Data Fig. 7). The widths of the fitted Gaussian correspond to two orthogonal diffusion components, from which we derived the associated diffusion coefficients.

The diffusion coefficients of the walkers are in the range of 1 – 2 × 10^3^ nm^2^/s (Extended Data Table 4), depending on the type of walker and track. These values are at least 10^4^ times lower compared to a protein of similar mass diffusing in water^19^. The combination of rapid diffusion in solution and pronounced anisotropy strongly supports a processive mode of movement along the fibres. Once dissociated, a walker would diffuse so quickly that it would vanish from the field of view. The likelihood of re-binding through repeated encounters (hop-diffusion) is minimal, as the three-dimensional aqueous volume dwarfs the two-dimensional track surface. Moreover, hop-diffusion would be insensitive to track spacing and isotropic in character. Measurements of a single foot binding and unbinding along the tracks show little anisotropy and significantly shorter dwell times. For comparison, kinesin-1 on microtubules in the absence of ATP^20^, diffuses only about tenfold faster than our walkers. Therefore, we focus on the largest principal component of the diffusion tensor, which captures motion along the track (Fig. 4a, right).

No binding was observed between the walkers and the track with the shortest footholds (TRACK-P3SN_3h-mNG). We measured the diffusion of walkers with varying numbers of feet along the remaining tracks (TRACK-P3SN_4h-mNG and P3SN_3.5h-mNG, Fig 4b, Extended Data Table 4). A principal trend emerged: the walkers with a more feet moved faster, while the binding affinity of the footholds did not have a significant effect. The average dwell time was shorter on tracks with weaker P3SN_3.5h feet (Extended Data Fig. 9, Extended Data Table 4).

The increase in diffusion rate might correspond to a higher local concentration of feet in higher-order oligomers, which causes an increase in the rate of association while the rate of dissociation remains unchanged^21^. Our observations agree with the results of simulations of the movement of multivalent ligands in two-dimensional space, which show that the diffusion rate increases with valency^22^.

To gain further insight into the correlation between affinity and diffusion rate we characterized the kinetics of single P3SN_4h, P3SN_3.5h and P4SN_4h feet using biolayer interferometry (BLI) and isothermal titration calorimetry (ITC; Extended Data Fig. 10, Extended Data Table 8). Truncating the P3SN coiled-coil by half a heptad reduced the association rate by approximately 23-fold and increased the dissociation rate 3.6-fold.

Based on single-foot kinetics, we anticipated a larger difference in diffusion rates between TRACK-P3SN_3.5h-mNG and TRACK-P3SN_4h-mNG. We performed BLI with single footholds immobilized on the chip and walkers as analytes (Extended Data Fig. 10, Extended Data Table 9). Even with a low density of footholds, where we assume the walker can only bind a single foothold, the difference in association rates between walkers on footholds P3SN_3.5h and P3SN_4h was less pronounced (∼ 3.6-fold) compared to that of single feet on the two footholds (∼ 23-fold). This valency effect could explain why the difference in diffusion between tracks with different footholds is smaller than would be expected from the affinity of single feet. The dwell time (Extended Data Fig. 9, Extended Data Table 4) of ∼ 70 seconds on P3SN_4h footholds is much larger than the half-lifetime of single feet (∼ 14 s).

To assess whether the mNeonGreen that is attached to the tracks influences walker mobility, we measured the diffusion of WALKER-C4 and WALKER-C8 on tracks lacking the fluorescent protein (Fig. 4c, Extended Data Table 6). Diffusion rates were comparable on tracks with and without mNeonGreen, indicating that the presence of flexibly tethered fusions does not measurably affect walker movement once bound to the track.

Next, we evaluated the effect of temperature, viscosity and salt concentration on movement of WALKER-C8 on tracks lacking mNeonGreen (Fig. 4d, Extended Data Table 6). While 20 % sucrose (1.695 cP at 25 °C) did not slow down the walker, its diffusion rate on both tracks dropped approximately 2-fold at 50 % sucrose (12.4 cP at 25 °C). The diffusion rate also dropped approximately 1.5-fold at 500 mM, most likely due to a decrease in affinity between the feet and the footholds. Conversely, when temperature was increased from 25 °C to 45 °C, their diffusion rates increased approximately 1.5-fold.

## Discussion

Here we report to the best of our knowledge the first fully *de novo* designed random protein walker. This system represents a milestone in the level of structural and functional complexity that can be achieved through protein design.

We engineered micron-scale fibres of large diameter and developed a versatile method (fibre H-fuse) to rigidly attach arbitrary protein domains to fibres based on secondary structure overlap. Cryo-EM analysis confirmed that the protomer structure closely matches the design model. We systematically tested a series of heterodimers for the ability to enable fibre growth and identified the most stable and reversible pairs. Building on this foundation, we designed walkers with varying valencies (i.e. differing numbers of feet) and showed that helical repeat proteins can be used to precisely tune inter-foot spacing.

To quantify the effect of valency and binding affinity, we performed extensive single-molecule tracking. Surprisingly, we find that walkers with a greater number of feet move faster. Based on the slow and anisotropic diffusion we conclude that we are observing diffusion along the fibres instead of a series of binding and unbinding events. We characterized the binding affinity and kinetics of both single feet and complete walkers and demonstrate that there is a valency effect even when the walkers bind to a single foothold. The number and binding affinity of feet, together with physical conditions, provide a powerful approach to fine-tune the system’s behaviour.

The walker operates without the need for specialized reagents or crosslinking chemicals, making it genetically encodable and possible to introduce into living cells using transfection. Because it is expected to be orthogonal to endogenous transport pathways based on actin and microtubule filaments, it may also open avenues for *in vivo* applications. Our *de novo* designed tuneable walker platform provides a general scaffold for building powered, programmable molecular machines moving forward.

## Acknowledgements

We thank L. Goldschmidt and K. VanWormer, for maintaining the computational and wet laboratory resources at the Institute for Protein Design.

This work was supported by the following funding sources: Slovenian research and Innovation agency N1-0323, BI-US/24-26-040, P4-0176 and European Union’s Horizon 2020: CC-LEGO 792305 to A.L., the Howard Hughes Medical Institute to H.S. and D.B., The Audacious Project at the Institute for Protein Design to H.S. and D.B., The Open Philanthropy Project Improving Protein Design Fund to Y.H. and D.B., Washington Research Foundation to J.W.

## Author Contributions

D.B. and A.L. directed the work, designed the research and led the project. H.S. designed and tested the HB67 fibre, including negative stain screening. A.L., with help from H.S. and Y.H, designed and tested the HA13 fibre. A.L. and Y.H. developed the fibre H-fuse protocol. A.L. designed and tested the tracks and walkers and screened heterodimers for binding. J.W. and J.D. helped with single molecule fluorescence tracking. E.L. and J.K determined the cryo-EM structure of HA13. T.S., E.R. and F.O. determined the cryo-EM structures of walkers. F.O. helped with modelling figure preparation. L.U. performed protein expression, refined the fibre formation protocol, performed BLI measurements, MP measurements, negative stain screening of tracks and all single molecule tracking experiments. E.R and L.U. performed ITC. A.L. wrote the tracking pipeline with help from L.U., J.D. and F.O. L.U and A.L. wrote the initial version of the manuscript. All authors reviewed and commented on the manuscript.

## Data availability

The structure of the track was deposited in the Protein Data Base (https://www.rcsb.org/) with the ID 9ZWG. Cryo-EM densities were deposited into the Electron Microscopy Data Bank (FIBRE-HA13: EMD-74908, WALKER-C4: EMD-54914, WALKER-C4b: EMD-54917, WALKER-C6: EMD-54911 and WALKER-C8: EMD-54912). ITC and BLI experiments were deposited to the Molecular Biophysics Database (https://mbdb-data.org/) (P4SN_4h vs P3SN_4h: ha172-vrw73 (BLI), bzrds-gmv73 (ITC), P4SN_4h vs P3SN_3.5h: y9h1c-mbk68 (BLI), qbtr9-ygc45 (ITC), P3SN_4h vs WALKER-C4: qbqwt-qjz12, P3SN_4h vs WALKER-C5: z74mj-fk011, P3SN_4h vs WALKER-C6: saf7x-cn219, P3SN_4h vs WALKER-C6b: sx065-bct52, P3SN_4h vs WALKER-C8: 25gan-csj07, P3SN_3.5h vs WALKER-C4: qe9tz-hax34, P3SN_3.5h vs WALKER-C5: 8c98z-jsr26, P3SN_3.5h vs WALKER-C6: k1×85-mmn68, P3SN_3.5h vs WALKER-C6b: jjpvk-ffx87, P3SN_3.5h vs WALKER-C8: 40xnq-76w56, P3SN_4h vs WALKER-C8, 10 nM load: ncmdg-zyt36, P3SN_4h vs WALKER-C8, 5 nM load: hmk82-3cz33, P3SN_4h vs WALKER-C8, 1 nM load: 0zeka-05z36, P3SN_3.5h vs WALKER-C8, 10 nM load: cyjzd-7e633, P3SN_3.5h vs WALKER-C8, 5 nM load: g8p0v-9at92). Models of walkers and fibres will be available on Zenodo (https://zenodo.org/) at publication time.

All other data are available in the main text or as Extended Data.

## Code availability

Code for the fibre H-fuse will be available on GitHub at publication time. Code for analysis of single molecule trajectories is available on GitHub at https://github.com/ajasja/walker_tracker and code for aligning TIRF and IRM channels is available at https://github.com/ajasja/LUMICKS_image_alignment.

## Extended Data Figures

**Extended Data Fig. 1:**
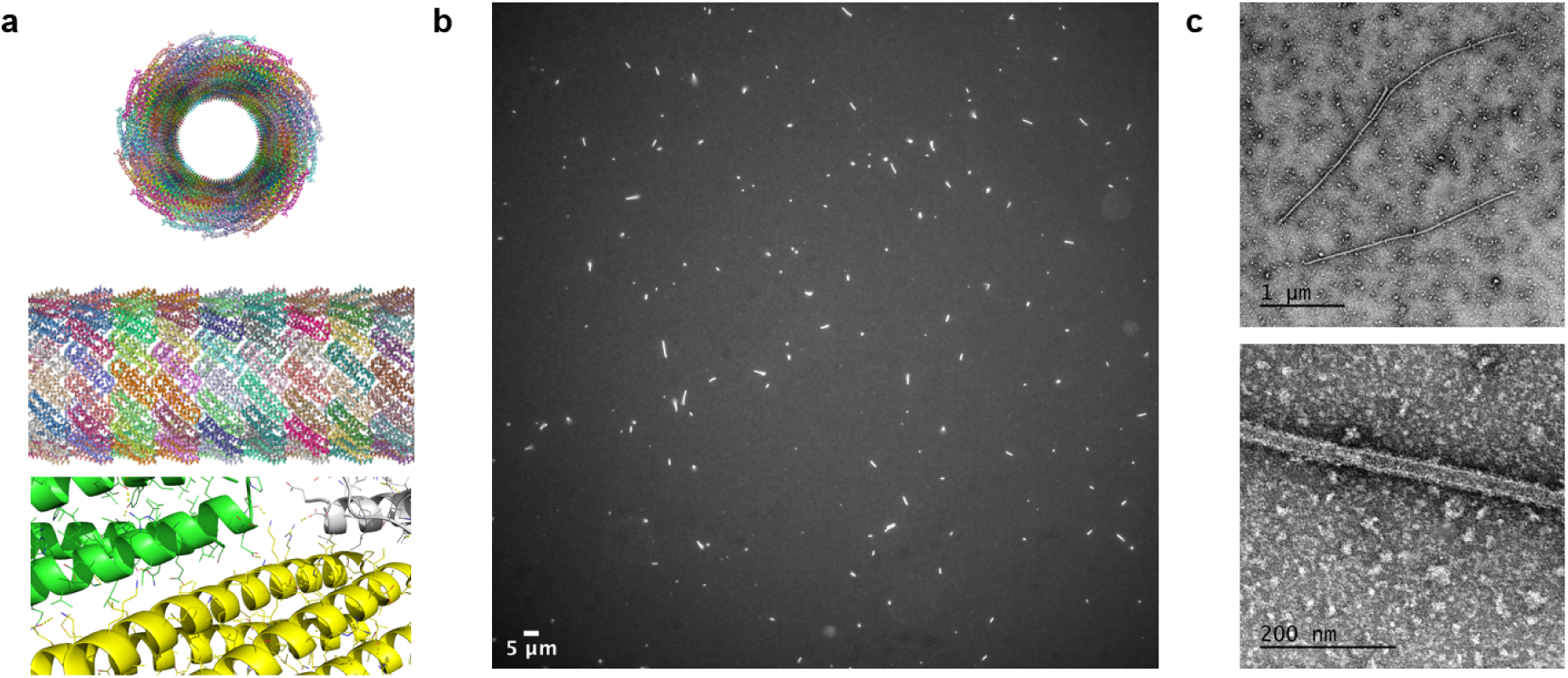
Design and characterisation of the HB67 large diameter fibre. **(a)** Computational design model of HB67, shown as top view (upper), side view (middle), and close-up of the inter-helical interface (lower). **(b)** Representative fluorescence micrograph of HB67 fibres labelled with GFP (scale bar, 5 μm). **(c)** Representative negative-stain electron micrographs of HB67 fibres, illustrating overall morphology (upper; scale bar, 1 μm) and higher-magnification view of fibre architecture (lower; scale bar, 200 nm).

**Extended Data Fig. 2:**
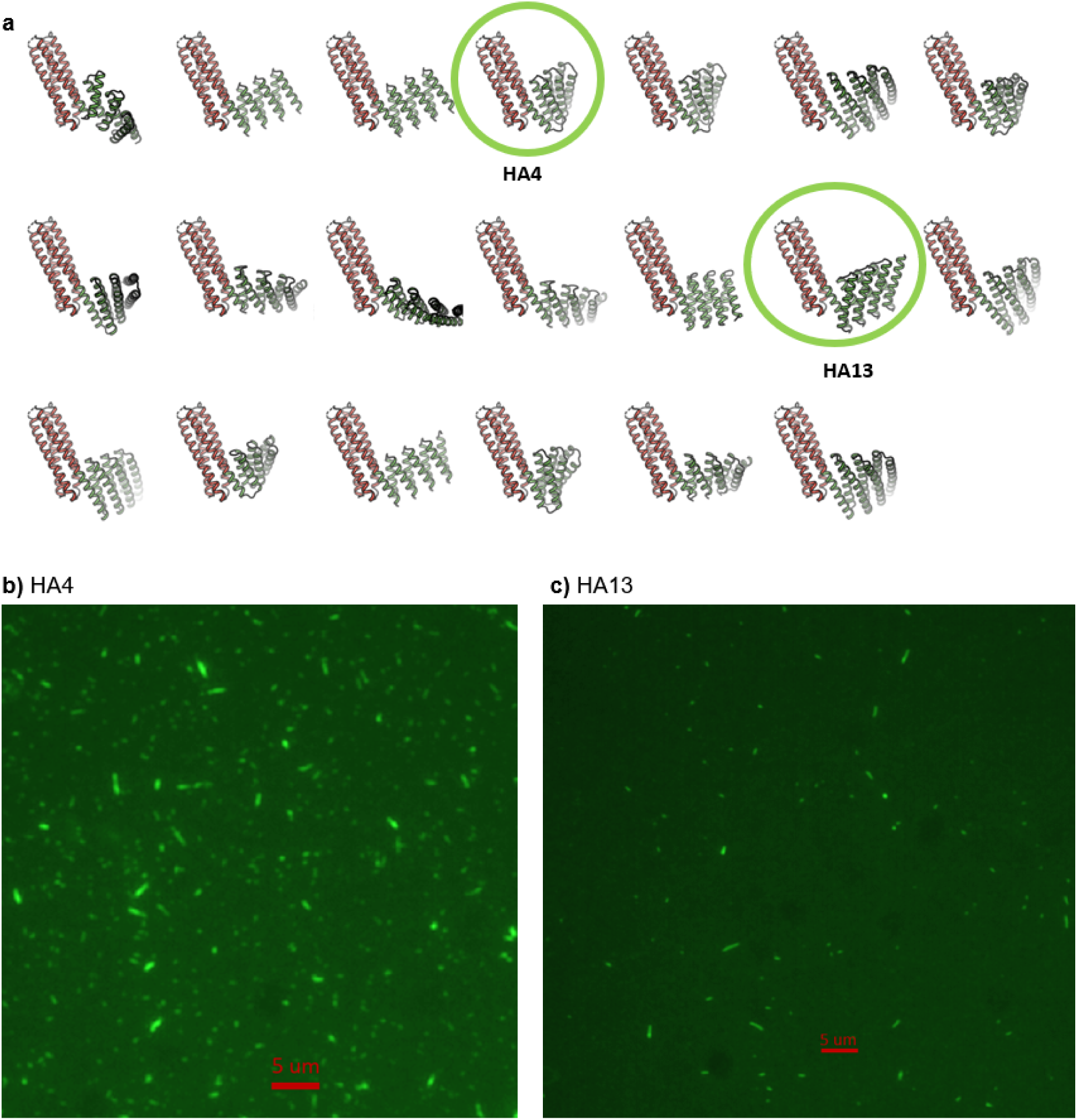
Design and characterization of HB67–DHR protein fusions. **(a)** Computational models of HB67 fused to designed helical repeat (DHR) proteins. HB67 fibres are shown in green and DHRs in red; selected designs HA4 and HA13 are highlighted. **(b)** Fluorescence microscopy image of GFP-tagged HA4 fibres. Scale bar, 5 µm. **(c)** Fluorescence microscopy image of GFP-tagged HA13 fibres. Scale bar, 5 µm.

**Extended Data Fig. 3:**
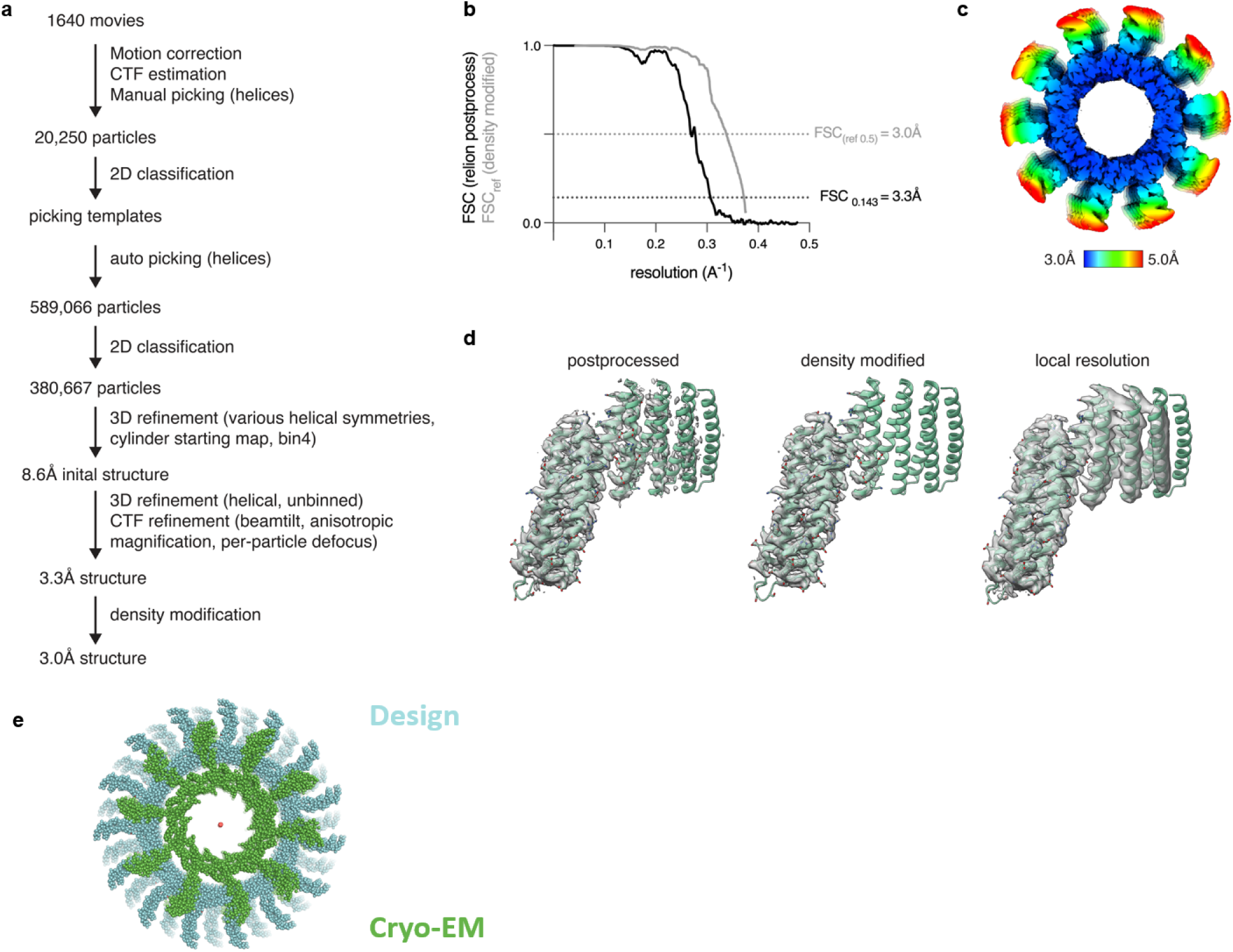
Details of the track structure and comparison to design model. **(a)** CryoEM processing workflow for HA13. **(b)** FSC curves for the HA13 cryoEM structure. The half-map FSC curve from relion postprocessing (black) and FSCref curve after density modification (grey) and corresponding resolution estimates are shown. **(c)** HA13 cryoEM structure colored by local resolution. **(d)** Atomic model of a single HA13 protomer together with maps from Relion postprocessing, density modification, or filtered according to local resolution. **(e)** Comparison of design model to Cryo-EM structure.

**Extended Data Fig. 4:**
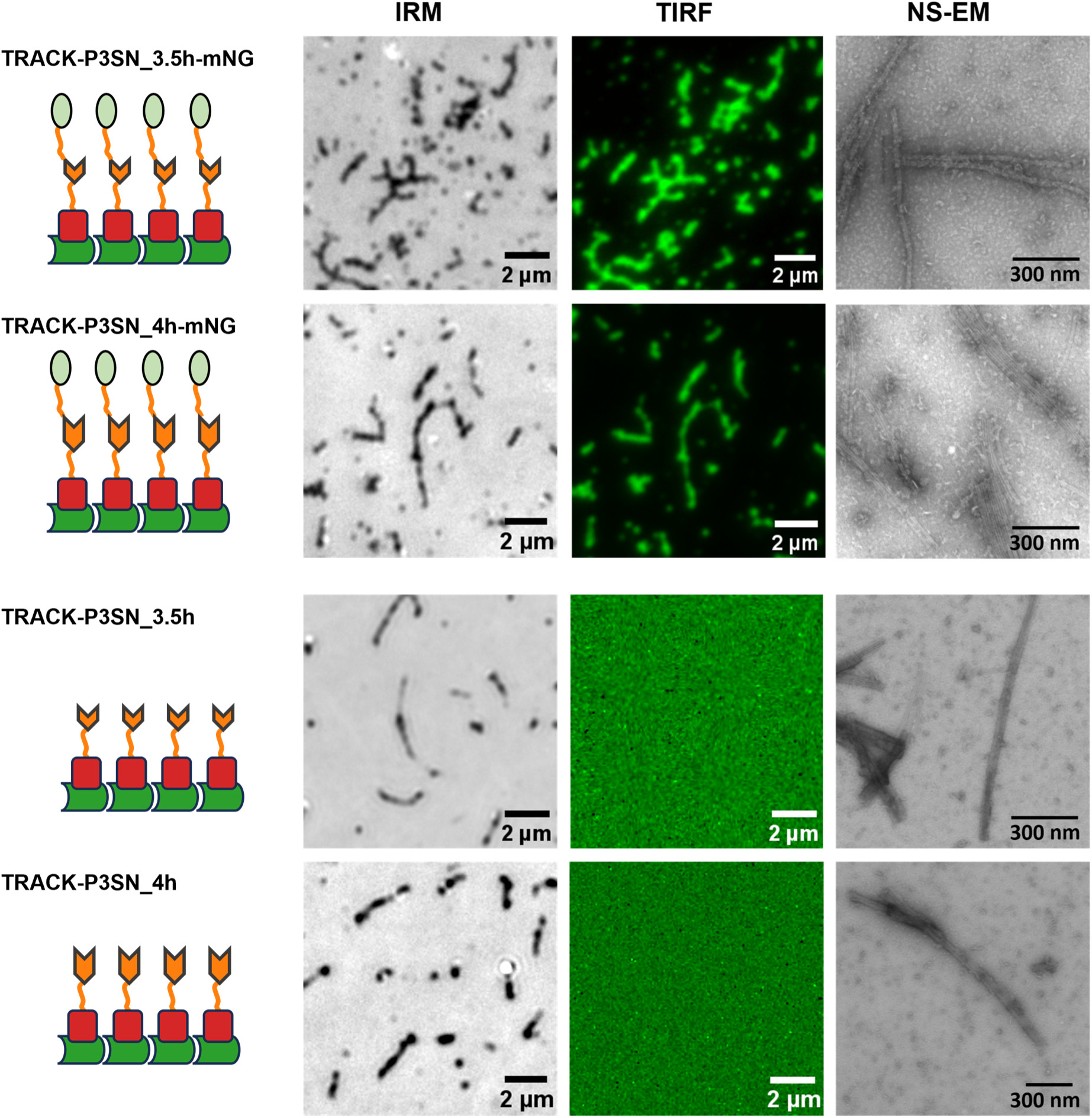
Experimental validation of the assembly of tracks used in diffusion experiments. Left, schematic representation of the designed fibres, showing the fibre backbone (dark green), DHR modules (red), footholds (orange), and mNeonGreen (mNG, light green). Representative images of the assembled structures obtained by interference reflection microscopy (IRM; scale bars, 2 µm), total internal reflection fluorescence microscopy (TIRF; scale bars, 2 µm), and negative-stain electron microscopy (NS-EM; scale bars, 300 nm) are shown from left to right.

**Extended Data Fig. 5:**
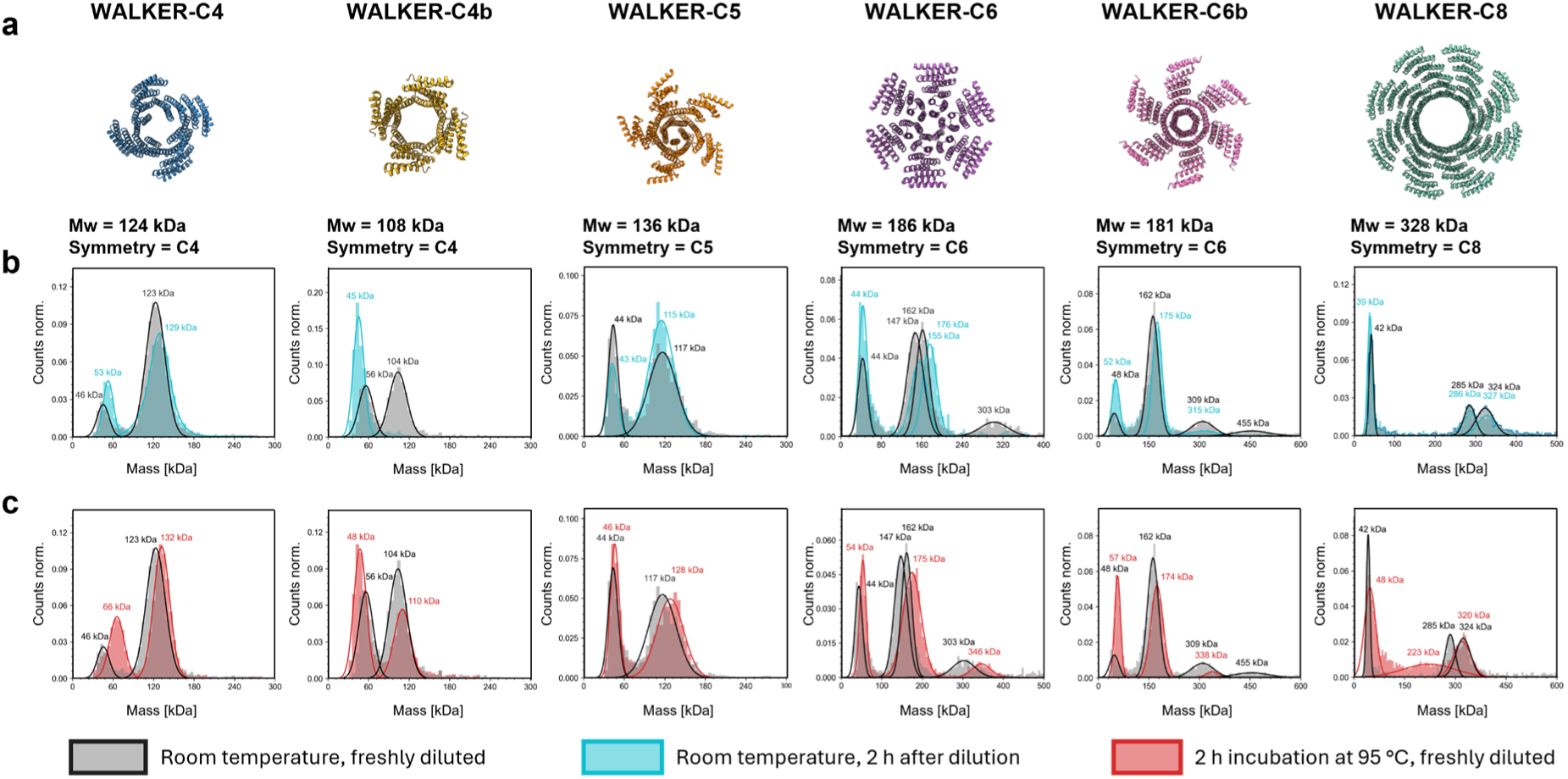
Oligomeric state of designed walkers assessed by mass photometry after prolonged incubation at low nanomolar concentrations and increased temperature. **(a)** Computational models of the designed walker oligomers, labelled according to their symmetry (e.g., WALKER-C4, -C5, -C8), with corresponding theoretical molecular weights (Mw) indicated. **(b)** Comparison of molecular weight distribution of freshly diluted walkers (grey) and walkers incubated at low nanomolar concentrations for 2 hours (cyan) both at room temperature. Five designs (WALKER_C4, WALKER_C5, WALKER_C6, WALKER_C6b, and WALKER_C8) maintained their expected oligomeric assemblies after prolonged incubation at low nanomolar concentrations. WALKER_C4b dissociated completely under the same conditions and was excluded from subsequent diffusion assays. **(c)** Comparison of molecular weight distribution of walkers incubated at room temperature (grey) and walkers incubated at 95 °C for 2 hours (red) and then diluted for MP. All designs retained their oligomeric state after incubation at 95 °C at high concentration for two hours.

**Extended Data Fig. 6:**
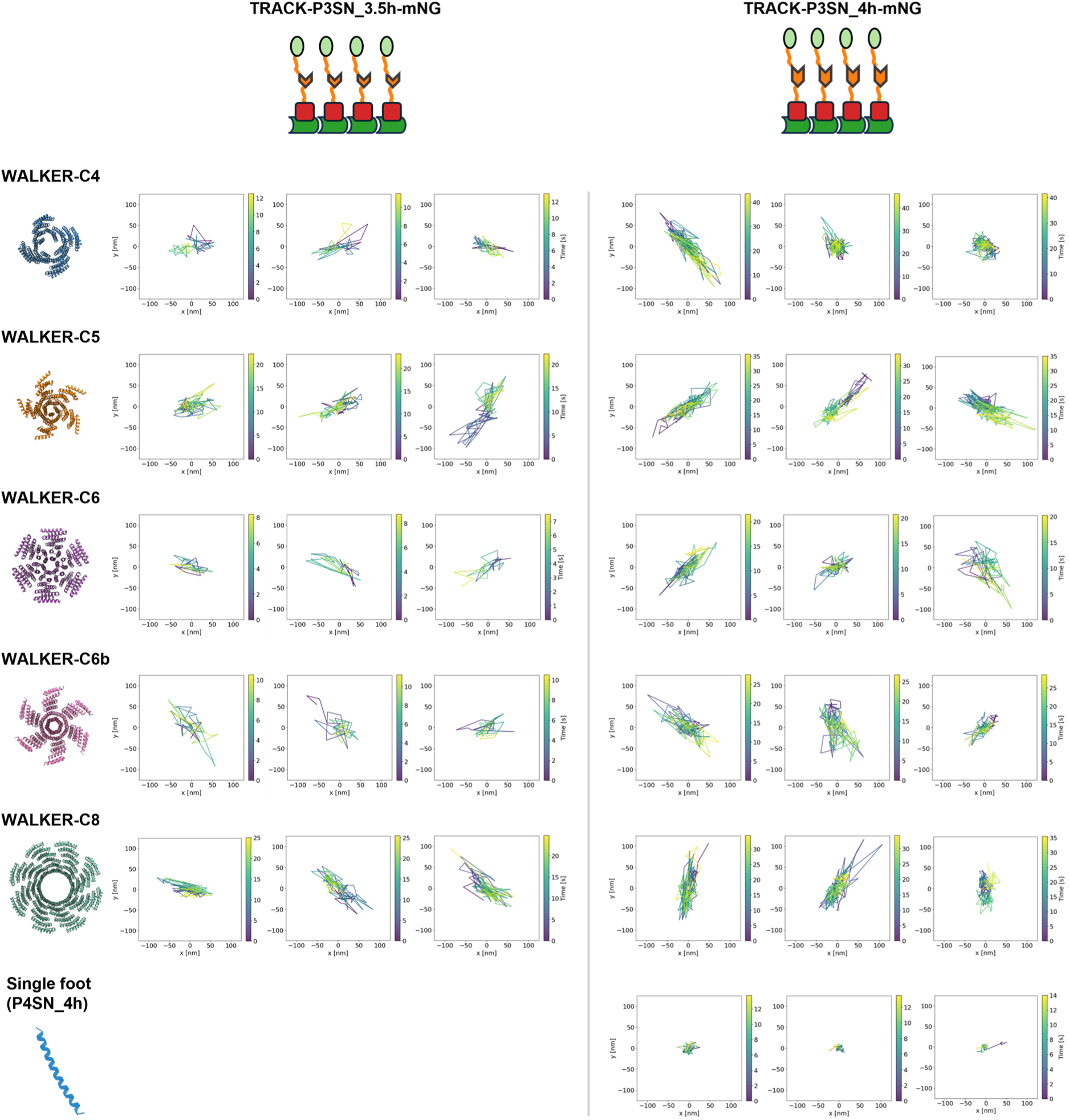
Representative trajectories of designed protein walkers on different tracks. For each walker–track combination, three representative single-walker trajectories are shown. Each plot corresponds to the trajectory of one walker, with position recorded over time. Position is in nanometres and colours (form purple to yellow) indicate the time from the beginning of the tracking in seconds.

**Extended Data Fig. 7:**
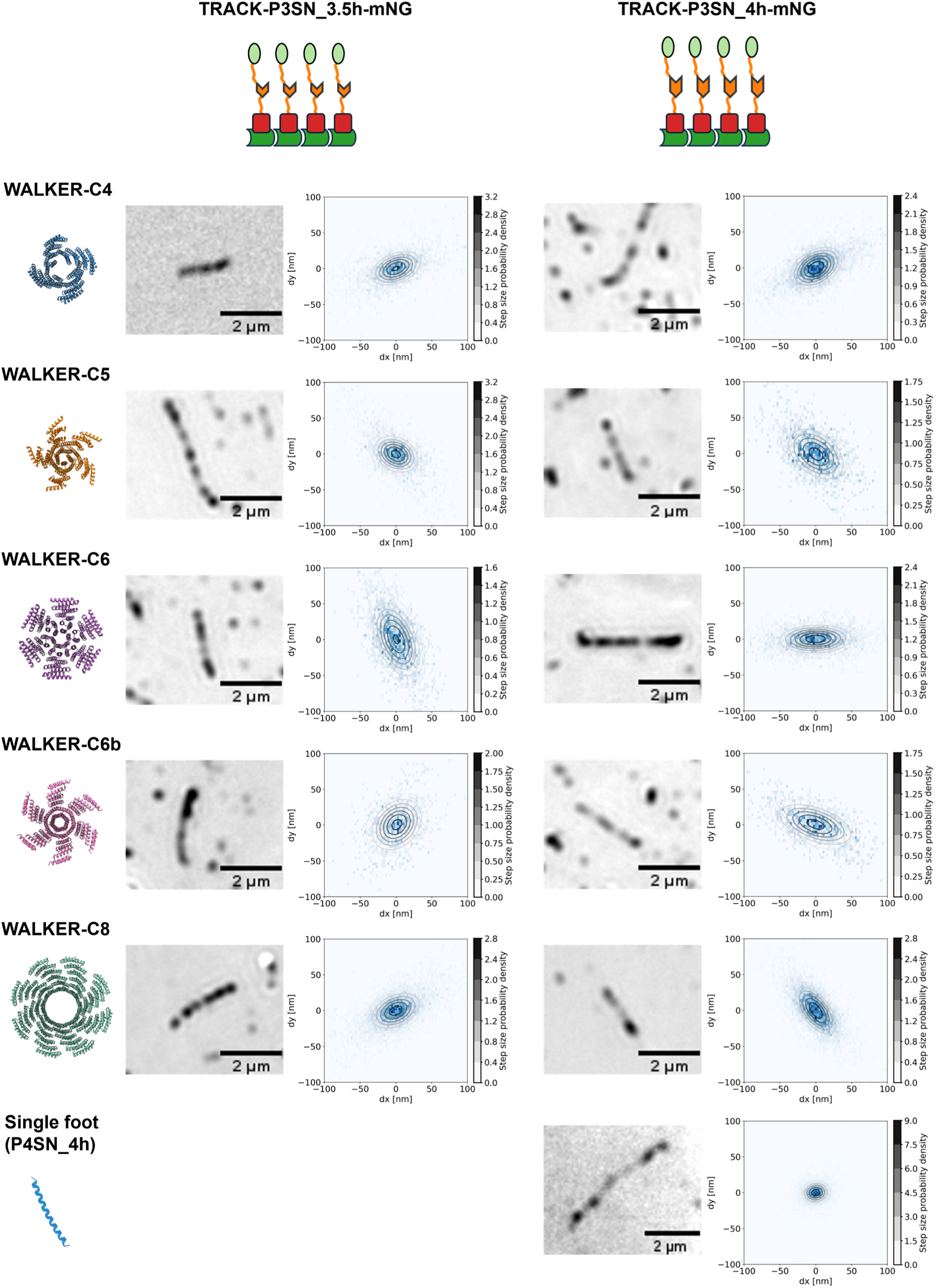
Examples of a two-dimensional distribution of steps sizes for each walker and track type used in diffusion experiments. Each histogram represents data collected from all walkers on the corresponding track, shown in the adjacent IRM micrograph to the left. The histograms are overlaid with fitted two-dimensional Gaussian contours representing probability density of step sizes. The elliptical shape of the probability density contours indicates anisotropic diffusion which is faster in the direction of the track. The rotation of the contours reflects the orientation of the track relative to the imaging axes.

**Extended Data Fig. 8:**
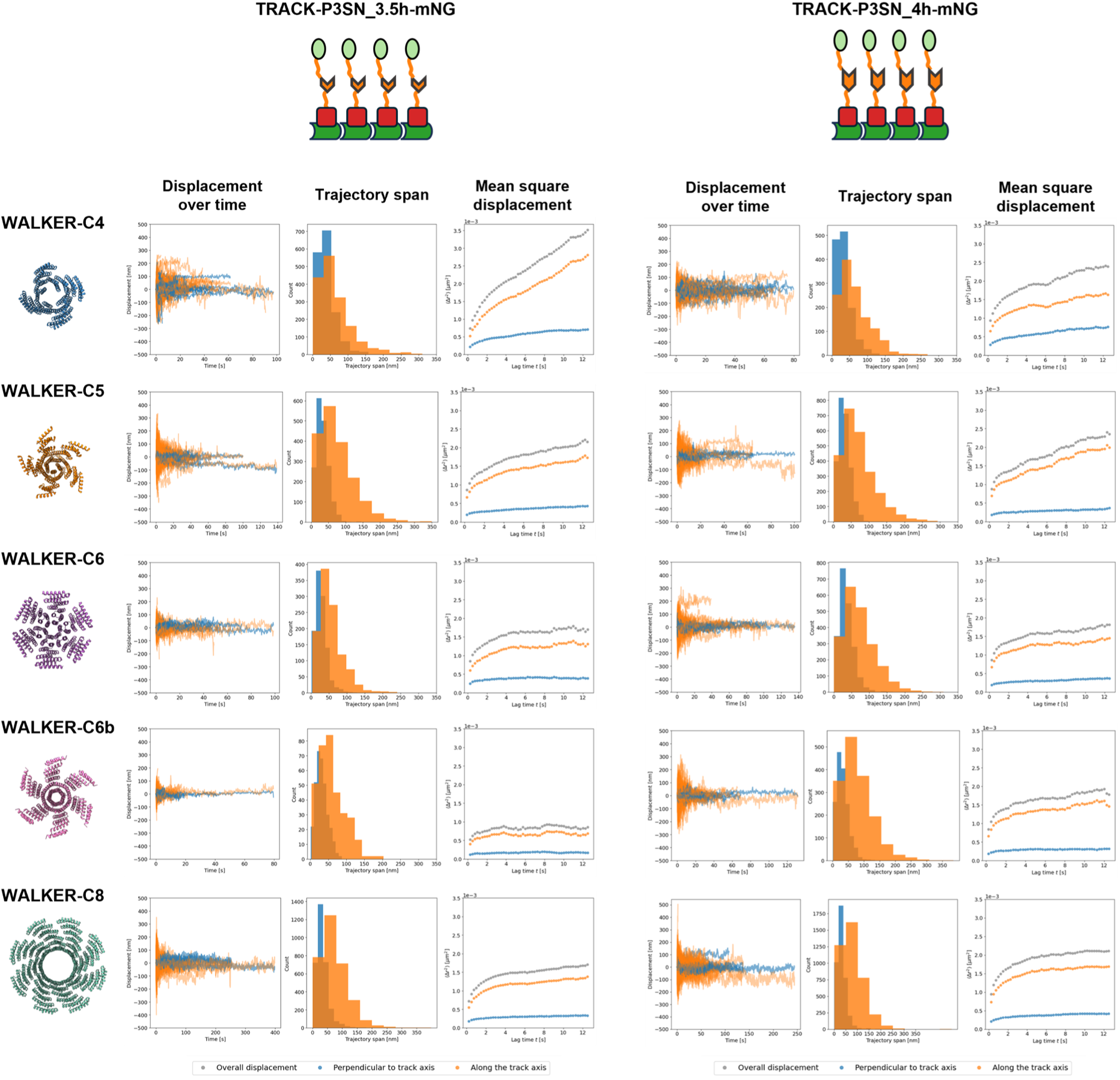
Analysis of the walkers’ trajectories along and perpendicular to the track axis. Displacement over time and trajectory span were greater and more variable along the track axis compared to the perpendicular direction, reflecting the inherent anisotropy of the track structure. The mean square displacement (MSD) analysis showed that the diffusion of the walkers is anomalous, which is reminiscent of many biological systems, where walker movement is constrained due to molecular crowding^23^ or the presence of obstacles^24,25^. Moreover, MSD analysis further confirmed the anisotropy of the system – while the movement is constrained in both dimensions, the confinement is more pronounced perpendicular to the track axis.

**Extended Data Fig. 9:**
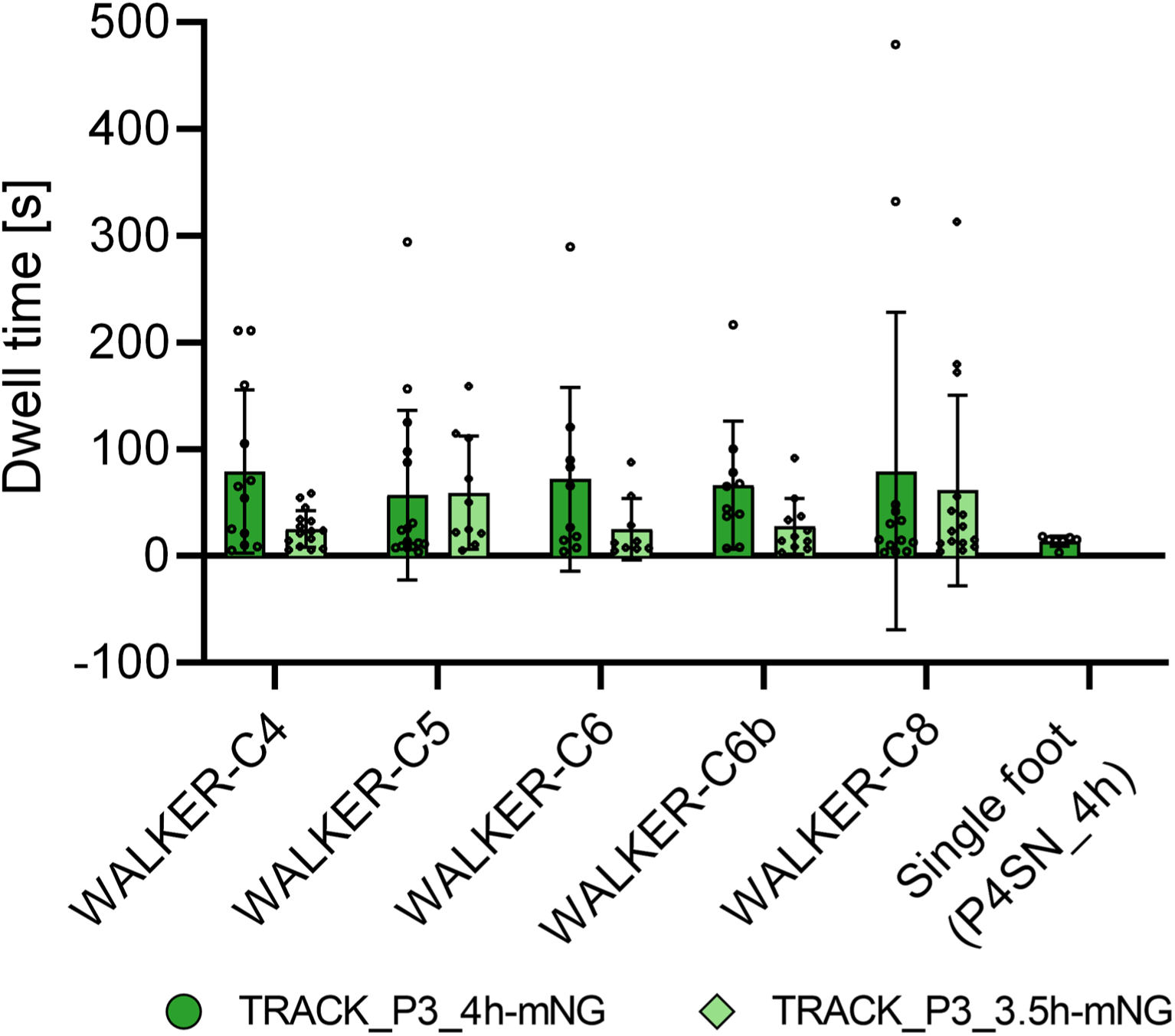
Average dwell times of walkers on both types of tracks (TRACK-P3SN_3.5h-mNG and TRACK-P3SN_4h-mNG). Dwell times were calculated using Trackpy, allowing a maximum gap of two frames to link walker localizations into trajectories. This accounts for fluorophore blinking between consecutive frames and yields more realistic estimates. On average, walkers exhibit longer dwell times—and thus greater processivity—on TRACK-P3SN_4h-mNG, consistent with the lower dissociation rates determined by BLI under sensor saturation conditions. Data represent mean ± s.d. from individual TIRF recordings.

**Extended Data Fig. 10:**
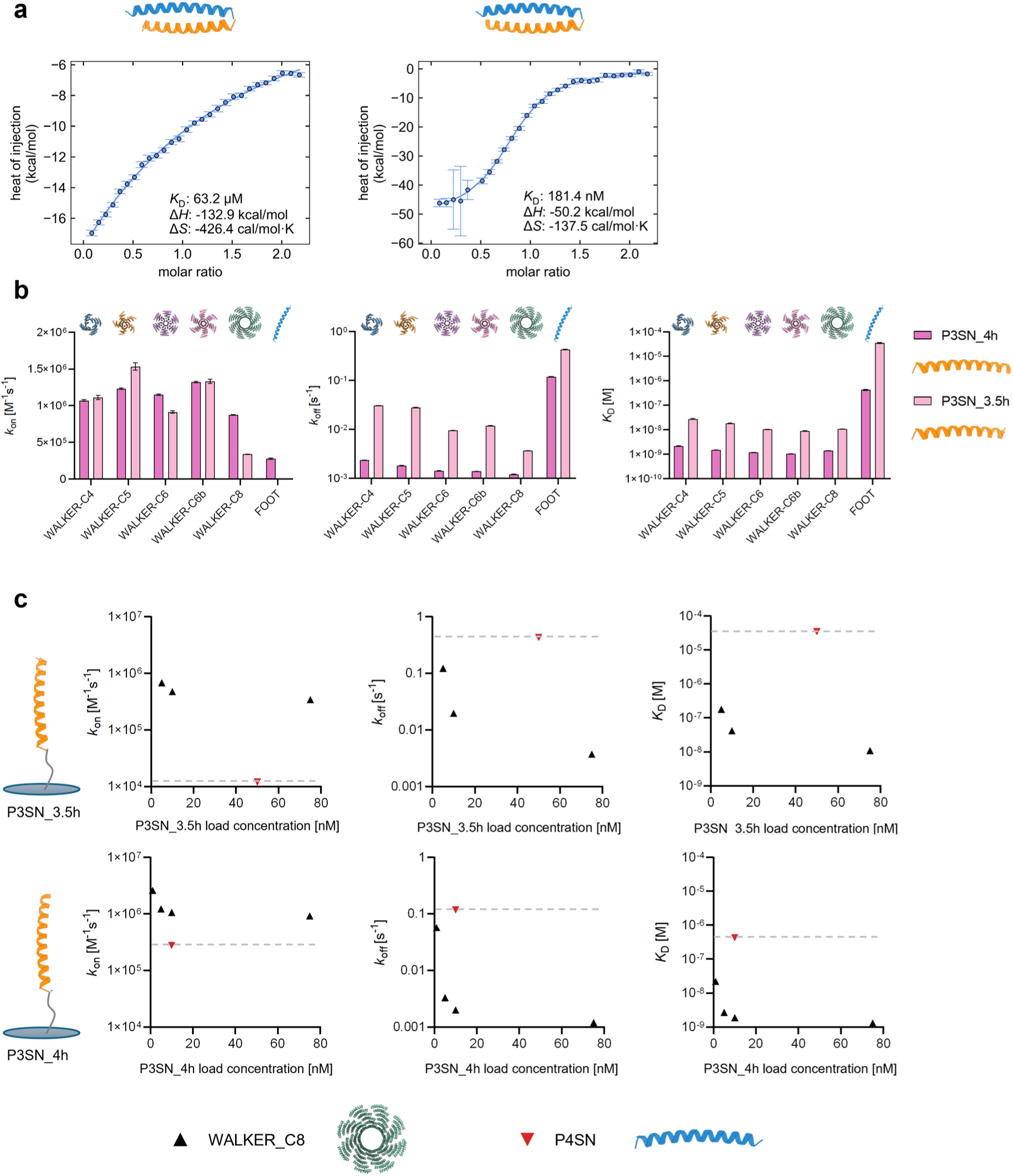
Kinetic and thermodynamic analysis of the interaction between walker feet and track footholds. (**a**) Isothermal titration (ITC) of peptide P4SN with peptides P3SN_3.5h (left) and P3SN_4h (right). (**b**) Kinetic parameters (*k*_on_, *k*_off_, *K*_D_) for walker and s foot binding to P3SN_3.5h and P3SN_4h, measured by BLI. Footholds were immobilized on the sensor to the point of saturation; walkers and feet served as analytes. The average spacing between footholds on the sensor surface is ∼ 17 nm^26^, allowing for multivalent attachment of walkers. (**c**) Dependence of kinetic parameters on foothold loading concentration. WALKER-C8 was used as the analyte; P3SN_3.5h and P3SN_4h served as ligands. The grey dashed line denotes the value of each kinetic parameter determined for single feet. At low loading (1 nM for P3SN_4h, 5 nM for P3SN_3.5h), walker dissociation rates approximate those of single feet, indicating a walker likely binds to a single foothold. We noticed that the measured kinetics depends on the density of the footholds. If the sensor is saturated with the footholds the difference in binding affinities is ∼ 10-fold smaller compared to single feet. At lower density of the footholds the dissociation rates approached that of an individual foothold, however association rates remained higher.

## Extended Data Tables

**Extended Data Table 1:** Fibres with various attached heterodimers screened for fibre formation. All heterodimeric constructs tested for fibre formation are listed. Fibre assembly was assessed by fluorescence microscopy. The “Fibre score” column reflects the observed morphology, on a scale from 0 to 3: 0, no fibres; 0.5, elongated spots (<1 µm); 1, short fibres (∼1 µm); 2, long fibres (2–5 µm); 3, very long fibres (>5 µm). Each entry also includes a brief description, reference, and the full construct sequence.

| Construct | Fibers | Short description | Reference | Sequence |
| --- | --- | --- | --- | --- |
| HA13LL_mNeonGreen | 1 | No heterodimers, positive control HB67 fibres with DHR | This study | MDSVEHLIKLLT |
| HA4LL_mNeonGreen | 2 | No heterodimers, positive control HB67 fibres with DHR | This study | MDSVEHLIKLLT |
| HA13LL_BW5A_mNeonGreen | 0 | Designed 4HB |  | MDSVEHLIKLLT |
| HA13LL_IL9B_mNeonGreen | 0 | Designed 4HB |  | MDSVEHLIKLLT |
| HA4LL_IL9B_mNeonGreen | 0 | Designed 4HB |  | MDSVEHLIKLLT |
| HA13LL_P3mSN_mNeonGreen | 2 | Designed coiled coil (TX -- number of truncated turns) | https://www. | MDSVEHLIKLLT |
| HA4LL_P3mSN_mNeonGreen | 1 | Designed coiled coil (TX -- number of truncated turns) | https://www. | MDSVEHLIKLLT |
| HA13LL_P3mSN_T1_mNeonGreen | 2 | Designed coiled coil (TX -- number of truncated turns) | https://www. | MDSVEHLIKLLT |
| HA4LL_P3mSN_T1_mNeonGreen | 2 | Designed coiled coil (TX -- number of truncated turns) | https://www. | MDSVEHLIKLLT |
| HA13LL_P3mSN_T2_mNeonGreen | 1 | Designed coiled coil (TX -- number of truncated turns) | https://www. | MDSVEHLIKLLT |
| HA13LL_P4mSN_mNeonGreen | 2 | Designed coiled coil (TX -- number of truncated turns) | https://www. | MDSVEHLIKLLT |
| HA4LL_P4mSN_mNeonGreen | 1 | Designed coiled coil (TX -- number of truncated turns) | https://www. | MDSVEHLIKLLT |
| HA4LL_P4mSN_T2_mNeonGree | 2 | Designed coiled coil (TX -- number of truncated turns) | https://www. | MDSVEHLIKLLT |
| HA4LL_P6mSN_mNeonGreen | 1 | Designed coiled coil (TX -- number of truncated turns) | https://www. | MDSVEHLIKLLT |
| HA13LL_T2B_A_mNeonGreen | 0.5 | Reversible designed four helix bundle mALb8 (TX -- number of truncated | https://www. | MDSVEHLIKLLT |
| HA4LL_T2B_A_mNeonGreen | 0.5 | Reversible designed four helix bundle mALb8 (TX -- number of truncated | https://www. | MDSVEHLIKLLT |
| HA4LL_T3A_A_mNeonGreen | 1 | Reversible designed four helix bundle mALb8 (TX -- number of truncated | https://www. | MDSVEHLIKLLT |
| HA13LL_T3B_A_mNeonGreen | 2 | Reversible designed four helix bundle mALb8 (TX -- number of truncated | https://www. | MDSVEHLIKLLT |
| HA4LL_T3B_A_mNeonGreen | 1 | Reversible designed four helix bundle mALb8 (TX -- number of truncated | https://www. | MDSVEHLIKLLT |
| FKBP_HA13LL_mNeonGreen | 3 | Rapamycin inducible heterodimer | https://pubs. | MDSVEHLIKLLT |
| FRB_HA13LL_mNeonGreen | 2 | Rapamycin inducible heterodimer | https://pubs. | MDSVEHLIKLLT |
| FRB_HA4LL_mNeonGreen | 1 | Rapamycin inducible heterodimer | https://pubs. | MDSVEHLIKLLT |
| LOV2_dark_HA13LL_mNeonGreen | 2 | Lov2 (light sensitive) mutant that is always in binding mode | https://www. | MDSVEHLIKLLT |
| LOV2_dark_HA4LL_mNeonGreen | 2 | Lov2 (light sensitive) mutant that is always in binding mode | https://www. | MDSVEHLIKLLT |
| LHD101A_HA13LL_mNeonGreen | 2 | Large designed heterodimer | https://www. | MDSVEHLIKLLT |
| LHD101A_HA4LL_mNeonGreen | 3 | Large designed heterodimer | https://www. | MDSVEHLIKLLT |
| LHD101B_HA13LL_mNeonGreen | 2 | Large designed heterodimer | https://www. | MDSVEHLIKLLT |
| LHD101B_HA4LL_mNeonGreen | 2 | Large designed heterodimer | https://www. | MDSVEHLIKLLT |
| LHD202A_HA13LL_mNeonGreen | 2 | Large designed heterodimer | https://www. | MDSVEHLIKLLT |
| LHD202A_HA4LL_mNeonGreen | 3 | Large designed heterodimer | https://www. | MDSVEHLIKLLT |
| HA13_LHD321A_mNeonGreen | 0 | Large designed heterodimer | https://www. | MDSVEHLIKLLT |
| HA4LL_LHD321A_mNeonGreen | 0.5 | Large designed heterodimer | https://www. | MDSVEHLIKLLT |
| ZDK1_HA13LL_mNeonGreen | 0 | Binds to Lov2 domain | https://www. | MDSVEHLIKLLT |
| ZDK1_HA4LL_mNeonGreen | 0 | Binds to Lov2 domain | https://www. | MDSVEHLIKLLT |
| ZDK2_HA13LL_mNeonGreen | 0.5 | Binds to Lov2 domain | https://www. | MDSVEHLIKLLT |
| ZDK2_HA4LL_mNeonGreen | 0.5 | Binds to Lov2 domain | https://www. | MDSVEHLIKLLT |
| AP3mS_HA13LL_mNeonGreen | 0 | Designed antiparallel coiled coil | https://www. | MDSVEHLIKLLT |
| AP3mS_HA4LL_mNeonGree | 0 | Designed antiparallel coiled coil | https://www. | MDSVEHLIKLLT |
| AP4_HA13LL_mNeonGreen | 0.5 | Designed antiparallel coiled coil | https://www. | MDSVEHLIKLLT |
| AP4_HA4LL_mNeonGreen | 1 | Designed antiparallel coiled coil | https://www. | MDSVEHLIKLLT |
| AP4mS_HA13LL_mNeonGreen | 3 | Designed antiparallel coiled coil | https://www. | MDSVEHLIKLLT |
| APHshSN_HA13LL_mNeonGreen | 1 | Designed antiparallel coiled coil | https://www. | MDSVEHLIKLLT |
| APHshSN_HA4LL_mNeonGreen | 1 | Designed antiparallel coiled coil | https://www. | MDSVEHLIKLLT |
| GCNshSN_HA13LL_mNeonGreen | 1 | Designed coiled coil (TX -- number of truncated turns) | https://www. | MDSVEHLIKLLT |
| GCNshSN_HA4LL_mNeonGreen | 2 | Designed coiled coil (TX -- number of truncated turns) | https://www. | MDSVEHLIKLLT |
| P5mSN_HA13LL_mNeonGreen | 3 | Designed coiled coil (TX -- number of truncated turns) | https://www. | MDSVEHLIKLLT |
| P5mSN_HA4LL_mNeonGreen | 2 | Designed coiled coil (TX -- number of truncated turns) | https://www. | MDSVEHLIKLLT |
| P6mSN_HA13LL_mNeonGreen | 0.5 | Designed coiled coil (TX -- number of truncated turns) | https://www. | MDSVEHLIKLLT |

**Extended Data Table 2:**
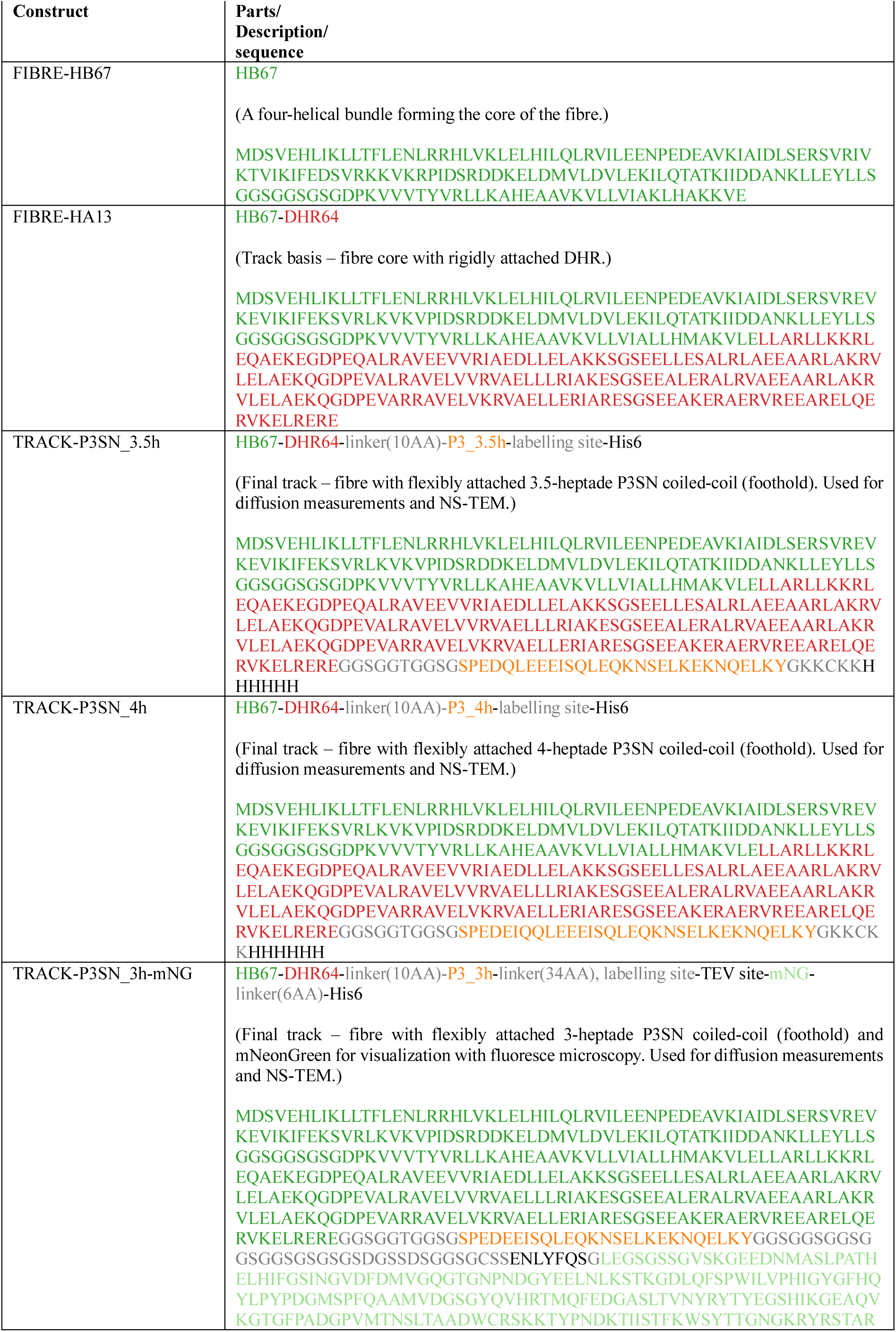
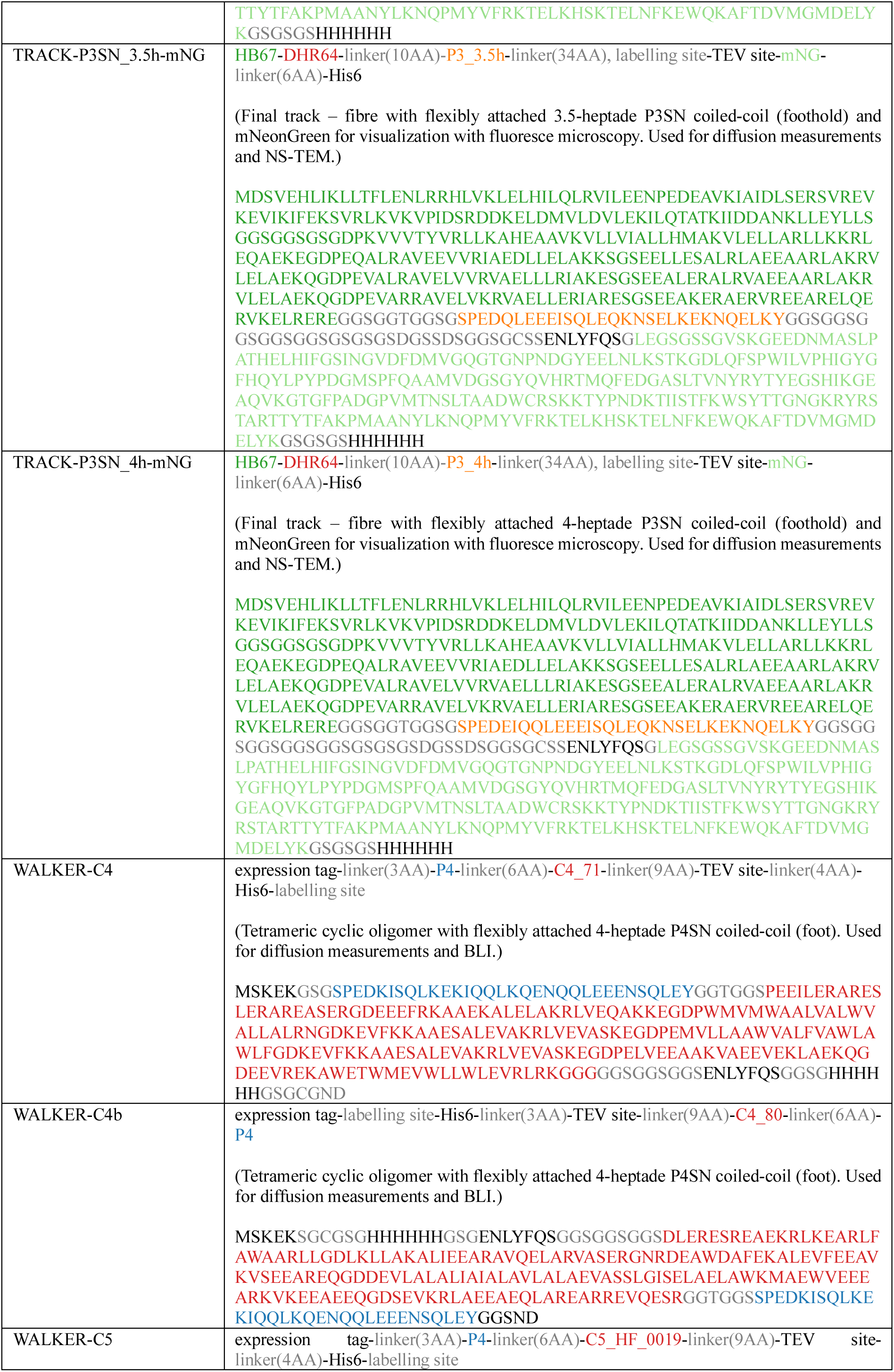
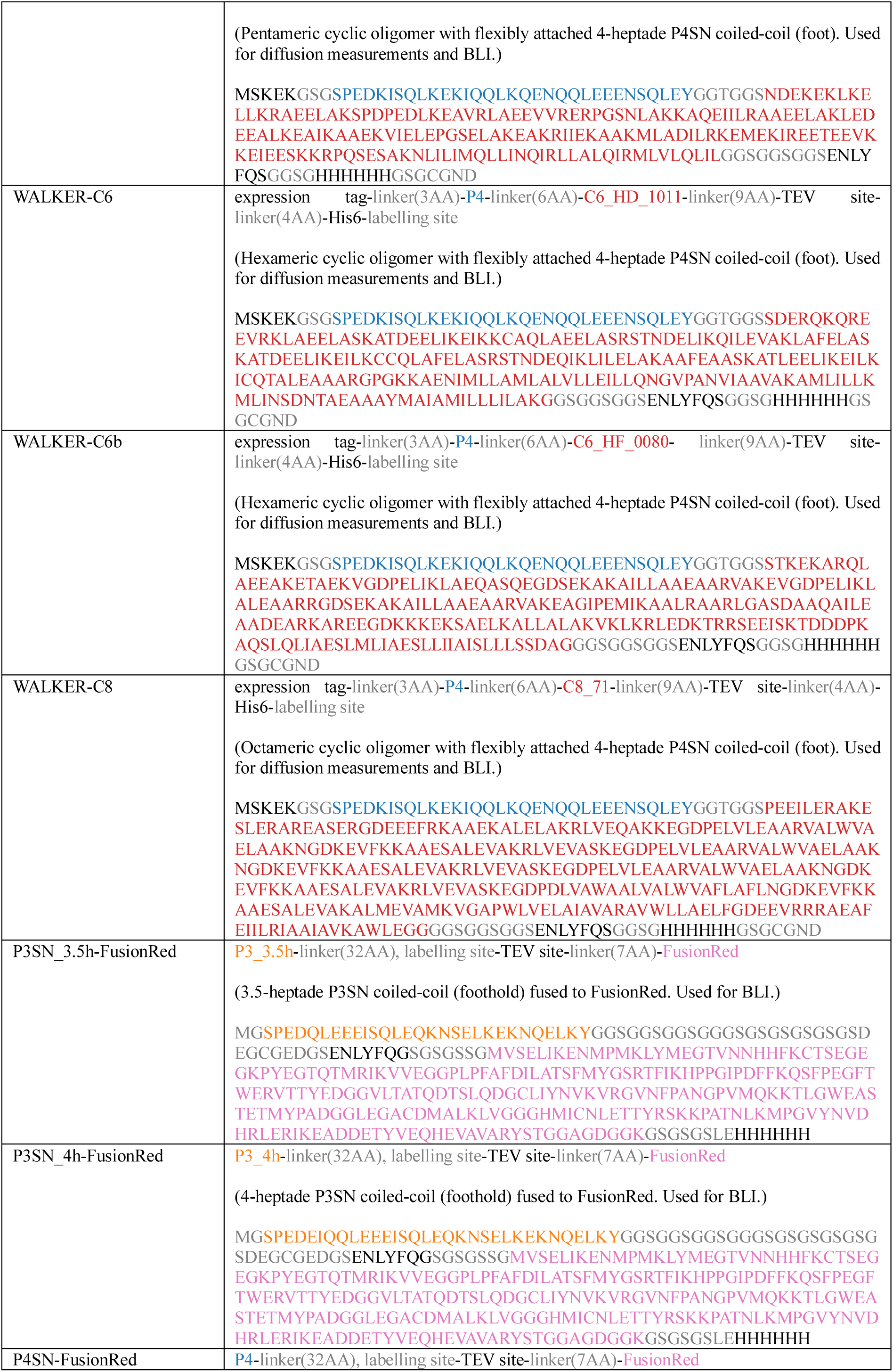
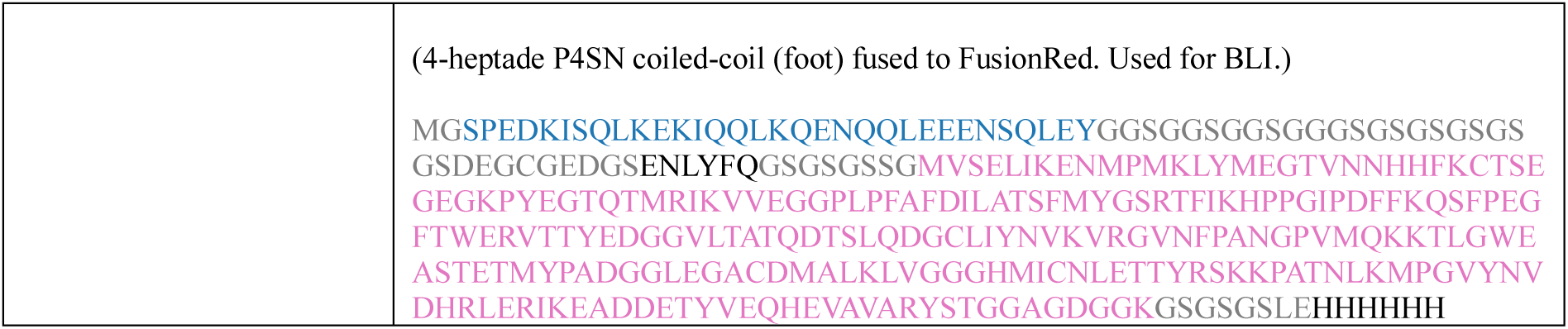
Names, descriptions and sequences of final constructs and their parts.

| Construct | Parts/<br>Description/<br>sequence |
| --- | --- |
| FIBRE-HB67 | **HB67**<br>(A four-helical bundle forming the core of the fibre.)<br>MDSVEHLIKLLTFLENLRRHLVKLELHILQLRVILEENPEDEAVKIAIDLSERSVRIV<br>KTVIKIFEDSVRKVKRPIDSRDDKELDMVLDVLEKILQTATKIIDDANKLLEYLLS<br>GGSGGSGSGDPKVVVTVYVRLKAHEAAVKVLLVIAKLHAKKVE |
| FIBRE-HA13 | **HB67-DHR64**<br>(Track basis – fibre core with rigidly attached DHR.)<br>MDSVEHLIKLLTFLENLRRHLVKLELHILQLRVILEENPEDEAVKIAIDLSERSVREV<br>KEVIKIFEKSVRLKVKVPIDSRDDKELDMVLDVLEKILQTATKIIDDANKLLEYLLS<br>GGSGGSGSGDPKVVVTVYVRLKAHEAAVKVLLVIALHMAKVLELLARLLKKRL<br>EQAEKEGDPEQALRAVEEVVRIAEDLLELAKKSGSEELLESALRLAEEAARLAKRV<br>LELAEKQGDPEVALRAVELVVRVAELLRIAKESGSEELERALRAVEEAARLAKR<br>VLELAEKQGDPEVARRAVELVKRVAELLERIAESGSEEAKEAERVREEARELQE<br>RVKELRERE |
| TRACK-P3SN_3.5h | **HB67-DHR64-linker(10AA)-P3_3.5h**-labelling site-His6<br>(Final track – fibre with flexibly attached 3.5-heptade P3SN coiled-coil (foothold). Used for diffusion measurements and NS-TEM.)<br>MDSVEHLIKLLTFLENLRRHLVKLELHILQLRVILEENPEDEAVKIAIDLSERSVREV<br>KEVIKIFEKSVRLKVKVPIDSRDDKELDMVLDVLEKILQTATKIIDDANKLLEYLLS<br>GGSGGSGSGDPKVVVTVYVRLKAHEAAVKVLLVIALHMAKVLELLARLLKKRL<br>EQAEKEGDPEQALRAVEEVVRIAEDLLELAKKSGSEELLESALRLAEEAARLAKRV<br>LELAEKQGDPEVALRAVELVVRVAELLRIAKESGSEELERALRAVEEAARLAKR<br>VLELAEKQGDPEVARRAVELVKRVAELLERIAESGSEEAKEAERVREEARELQE<br>RVKELRERE<br>GGSGGTGGSGSPEDQLEEEISQLEQKNSSELKEKNQELKYGKKCKKH<br>HHHHH |
| TRACK-P3SN_4h | **HB67-DHR64-linker(10AA)-P3_4h**-labelling site-His6<br>(Final track – fibre with flexibly attached 4-heptade P3SN coiled-coil (foothold). Used for diffusion measurements and NS-TEM.)<br>MDSVEHLIKLLTFLENLRRHLVKLELHILQLRVILEENPEDEAVKIAIDLSERSVREV<br>KEVIKIFEKSVRLKVKVPIDSRDDKELDMVLDVLEKILQTATKIIDDANKLLEYLLS<br>GGSGGSGSGDPKVVVTVYVRLKAHEAAVKVLLVIALHMAKVLELLARLLKKRL<br>EQAEKEGDPEQALRAVEEVVRIAEDLLELAKKSGSEELLESALRLAEEAARLAKRV<br>LELAEKQGDPEVALRAVELVVRVAELLRIAKESGSEELERALRAVEEAARLAKR<br>VLELAEKQGDPEVARRAVELVKRVAELLERIAESGSEEAKEAERVREEARELQE<br>RVKELRERE<br>GGSGGTGGSGSPEDEIQLEEEISQLEQKNSSELKEKNQELKYGKKCKK<br>KHHHHH |
| TRACK-P3SN_3h-mNG | **HB67-DHR64-linker(10AA)-P3_3h-linker(34AA)**, labelling site-TEV site-mNG-linker(6AA)-His6<br>(Final track – fibre with flexibly attached 3-heptade P3SN coiled-coil (foothold) and mNeonGreen for visualization with fluoresce microscopy. Used for diffusion measurements and NS-TEM.)<br>MDSVEHLIKLLTFLENLRRHLVKLELHILQLRVILEENPEDEAVKIAIDLSERSVREV<br>KEVIKIFEKSVRLKVKVPIDSRDDKELDMVLDVLEKILQTATKIIDDANKLLEYLLS<br>GGSGGSGSGDPKVVVTVYVRLKAHEAAVKVLLVIALHMAKVLELLARLLKKRL<br>EQAEKEGDPEQALRAVEEVVRIAEDLLELAKKSGSEELLESALRLAEEAARLAKRV<br>LELAEKQGDPEVALRAVELVVRVAELLRIAKESGSEELERALRAVEEAARLAKR<br>VLELAEKQGDPEVARRAVELVKRVAELLERIAESGSEEAKEAERVREEARELQE<br>RVKELRERE<br>GGSGGTGGSGSPEDEEISQLEQKNSSELKEKNQELKYGGSGGSGSGSG<br>GSGSGSGSGSDGSSDSGSGSCSSENLYFQSGLEGSGSSGVSKGEEDNMA SPLATH<br>ELHIFGSINGVDFDMVGQGTGNPNDGYEELNLKSTKGDLYFSPWILVPHIGYGFHQ<br>YLPYPDGMSPFQAAMVDGSGYQVHRTMQFEDGASLTVNYRYTYEGSHIKGEAQV<br>KGTGFPADGPVMTNSLTAADWCRSKKTYPNDKTIISTFKWSYTTGNGKRYRSTAR |
|  | TTYTFAKPMAANYLKNQPMYVFRKTELKHSKTELNFKEWQKAFTDVMGMDELYKGGSGSGSHHHHHH |
| TRACK-P3SN_3.5h-mNG | **HB67-DHR64**-linker(10AA)-**P3_3.5h**-linker(34AA), labelling site-TEV site-mNG-linker(6AA)-His6<br>(Final track – fibre with flexibly attached 3.5-heptade P3SN coiled-coil (foothold) and mNeonGreen for visualization with fluoresce microscopy. Used for diffusion measurements and NS-TEM.)<br>MDSVEHLIKLLTFLENLRRHLVKLELHILQLRVILEENPEDEAVKIAIDLSERSVREV<br>KEVIKIFEKSVRLKVKVPIDSRDDKELDMVLDVLEKILQTATKIIDDANKLLEYLLS<br>GGSGGGSGSDPKVVVTVYVRLKAHEAAVKVLLVIALHMAKVLELLARLLKKRL<br>EQAEKEGDPEQALRAVEEVVRIAEDLLELAKKSGSEELLESALRLAEEAARLAKRV<br>LELAEKQGDPEVALRAVELVVRVAELLRIAKESGSEELERARVAEEAARLAKR<br>VLELAEKQGDPEVARRAVELVKRVAELLERIAESGSEEAKEARAERVREEARELQE<br>RVKELREREGGSGGTGGSG**SPEDQLEEEISQLEQKNSSELKEKNQELKY**GGSGGGSG<br>GGSGGGSGSGSGSDGSSDSGGSGCSENLYFQSGLEGSGSSGVSKGEEDNMASLP<br>ATHELHIFGSINGVDFDMVGGGTGNPNDGYEELNLKSTKGDQLQFSPWILVPHIGY<br>FHQYLPYPDGMSPFQAAMVDGSGYQVHRTMQFEDGASLTVNRYTYEGSHIKGE<br>AQVKGTGFPADGPVMTNSLTAADWCERSKKTYPNDKTIISTFKWSYTTGNGKRYRS<br>TARTTYTFAKPMAANYLKNQPMYVFRKTELKHSKTELNFKEWQKAFTDVMGMD<br>ELYKGGSGSGSHHHHHH |
| TRACK-P3SN_4h-mNG | **HB67-DHR64**-linker(10AA)-**P3_4h**-linker(34AA), labelling site-TEV site-mNG-linker(6AA)-His6<br>(Final track – fibre with flexibly attached 4-heptade P3SN coiled-coil (foothold) and mNeonGreen for visualization with fluoresce microscopy. Used for diffusion measurements and NS-TEM.)<br>MDSVEHLIKLLTFLENLRRHLVKLELHILQLRVILEENPEDEAVKIAIDLSERSVREV<br>KEVIKIFEKSVRLKVKVPIDSRDDKELDMVLDVLEKILQTATKIIDDANKLLEYLLS<br>GGSGGGSGSDPKVVVTVYVRLKAHEAAVKVLLVIALHMAKVLELLARLLKKRL<br>EQAEKEGDPEQALRAVEEVVRIAEDLLELAKKSGSEELLESALRLAEEAARLAKRV<br>LELAEKQGDPEVALRAVELVVRVAELLRIAKESGSEELERARVAEEAARLAKR<br>VLELAEKQGDPEVARRAVELVKRVAELLERIAESGSEEAKEARAERVREEARELQE<br>RVKELREREGGSGGTGGSG**SPEDQLEEEISQLEQKNSSELKEKNQELKY**GGSGG<br>GGSGGGSGSGSGSDGSSDSGGSGCSENLYFQSGLEGSGSSGVSKGEEDNMAS<br>LPATHELHIFGSINGVDFDMVGGGTGNPNDGYEELNLKSTKGDQLQFSPWILVPHIG<br>YGFHQLPYPDGMSPFQAAMVDGSGYQVHRTMQFEDGASLTVNRYTYEGSHIK<br>GEAQVKGTGFPADGPVMTNSLTAADWCERSKKTYPNDKTIISTFKWSYTTGNGKRY<br>RSTARTTYTFAKPMAANYLKNQPMYVFRKTELKHSKTELNFKEWQKAFTDVMG<br>MDELYKGGSGSGSHHHHHH |
| WALKER-C4 | expression tag-linker(3AA)-**P4**-linker(6AA)-**C4_71**-linker(9AA)-TEV site-linker(4AA)-His6-labelling site<br>(Tetrameric cyclic oligomer with flexibly attached 4-heptade P4SN coiled-coil (foot). Used for diffusion measurements and BLI.)<br>MSKEKSGSG**SPEDKISQLKEKIQQLKQENQQLEEEENSQLEY**GGTGGSP**EEILERARES**<br>**LERAREASERGD**EEEEFRKAAEKALELAKRLVEQAKKEGDPWMVMWAALVALWV<br>ALLALRNGDKEVFKKAAESALEVAKRLVEVASKEGDPPEMVLLAAWVALFVAWLA<br>WLFGDKEVFKKAAESALEVAKRLVEVASKEGDPPELVAAKVAEEVEKLAQKQ<br>DEEVREKAWETWMEVWLLWLEVRLRKGGGGGGSGSGGSENLYFQSGSGSGHHHH<br>HHGSGCGND |
| WALKER-C4b | expression tag-labelling site-His6-linker(3AA)-TEV site-linker(9AA)-**C4_80**-linker(6AA)-**P4**<br>(Tetrameric cyclic oligomer with flexibly attached 4-heptade P4SN coiled-coil (foot). Used for diffusion measurements and BLI.)<br>MSKEKSGCGSGHHHHHHGSGENLYFQSGSGSGSGSD**LERESREAEKRLKEARLF**<br>**AWAARLLGDLKLLAKALIEEARAVQELARVASERGNRDEAWDAFEKALEVFEEAV**<br>**KVSEEA**REQDDEVLLALALIALAVLALAEVASSLGISELAELAWKMAEWVEE<br>**ARKVKEEAEEQGDSEVKRLAEAEQLAREARREVRQESR**GGTGGSP**EDKISQLKE**<br>**KIQQLKQENQQLEEEENSQLEY**GGSGND |
| WALKER-C5 | expression tag-linker(3AA)-**P4**-linker(6AA)-**C5_HF_0019**-linker(9AA)-TEV site-linker(4AA)-His6-labelling site |
|  | (Pentameric cyclic oligomer with flexibly attached 4-heptade P4SN coiled-coil (foot). Used for diffusion measurements and BLI.)<br>MSKEKGSGSPEDKISQLKEKIQQLKQENQQLEEEENSQLEYGGTGGSNDEKEKLKE<br>LLKRAEELAKSPDPEDLKEAVRLAEEVVRERPGSNLAKKAQEILRAAEELAKLED<br>EEALKEAIKAAEKVIELEPGSELAKEAKRIIEKAAKMLADILRKEMEKIREETEEVK<br>KEIEESKKRPQSESAKNLILIMQLLINQIRLLALQIRMLVLQLILGGSGSGSGSENLY<br>FQSGSGSGHHHHHHGSGCGND |
| WALKER-C6 | expression tag-linker(3AA)-P4-linker(6AA)-C6_HD_1011-linker(9AA)-TEV site-linker(4AA)-His6-labelling site<br>(Hexameric cyclic oligomer with flexibly attached 4-heptade P4SN coiled-coil (foot). Used for diffusion measurements and BLI.)<br>MSKEKGSGSPEDKISQLKEKIQQLKQENQQLEEEENSQLEYGGTGGSNDERQKQRE<br>EVRKLAEEELASKATDEELIKEIKKCAQLAEELASRSTNDELIQILEVAKLAFELAS<br>KATDEELIKEILKCCQLAFELASRSTNDEQIKLILELAKAAFEAASKATLEELIKEILK<br>ICQTALEAAARGPGKKAENIMLLAMLALVLEILLQNGVPANVIAAVAKAMLILLK<br>MLINSNTAEAAAYMAIAMILLILAKGSGSGSGSGSENLYFQSGSGSGHHHHHHGSG<br>GCGND |
| WALKER-C6b | expression tag-linker(3AA)-P4-linker(6AA)-C6_HF_0080-linker(9AA)-TEV site-linker(4AA)-His6-labelling site<br>(Hexameric cyclic oligomer with flexibly attached 4-heptade P4SN coiled-coil (foot). Used for diffusion measurements and BLI.)<br>MSKEKGSGSPEDKISQLKEKIQQLKQENQQLEEEENSQLEYGGTGGSSTKEKARQL<br>AEEAKETAEKVGDPELIKLAEQASQEGDSEKAKAILLAEEAARVAKEVGDPELIK<br>ALEAARRGDSEKAKAILLAEEAARVAKEAGIPEMIKAAALRAARLGASDAAQAILE<br>AADEARKAREEGDKKKEKS AELKALLALAKVKLKRLEDKTRRSEESKTDDEDPK<br>AQSLQLIAESLMLIAESLLIIAISLLSSDAGGSGSGSGSGSENLYFQSGSGSGHHHHHH<br>GSGCGND |
| WALKER-C8 | expression tag-linker(3AA)-P4-linker(6AA)-C8_71-linker(9AA)-TEV site-linker(4AA)-His6-labelling site<br>(Octameric cyclic oligomer with flexibly attached 4-heptade P4SN coiled-coil (foot). Used for diffusion measurements and BLI.)<br>MSKEKGSGSPEDKISQLKEKIQQLKQENQQLEEEENSQLEYGGTGGSPEEILERAKE<br>SLERAREASERGDEEEFRKAAEKALELAKRLVEQAKKEGDPELVLEAARVALWVA<br>ELAAKNGDKVEFKKAAESALEVAKRLVEVASKEGDPELVLEAARVALWVAELAAK<br>NGDKVEFKKAAESALEVAKRLVEVASKEGDPELVLEAARVALWVAELAAKNGDK<br>EVFKKAAESALEVAKRLVEVASKEGDPDLVAAALVALWVAFLNGDKVEFKK<br>AAESALEVAKALMEVAMKVGAPWLVELAIAVARAVWLLAELFGDEEVRRAEAF<br>EILRIAIAIAVKAWLEGGGSGSGSGSGSENLYFQSGSGSGHHHHHHGSGCGND |
| P3SN_3.5h-FusionRed | P3_3.5h-linker(32AA), labelling site-TEV site-linker(7AA)-FusionRed<br>(3.5-heptade P3SN coiled-coil (foothold) fused to FusionRed. Used for BLI.)<br>MGSPEDQLEEEISQLEQKNSSELKEKNQELKYGGSGSGSGSGSGSGSGSGSGSD<br>EGCGEDGSENLYFQSGSGSGSGMVSELIKENMPMKLYMEGTVNNHHFKCTSEGE<br>GKPYEGTQTMRIKVVEGGPLPFAFDILATSFMYGSRTFIKHPPGIPDFFKQSFPEGFT<br>WERVTTYEDGGVLTATQDTSQDGLIYNVVKVRGVNFPANGPVMQKKTGWEAS<br>TETMYPADGGLEGACDMALKLVGGGHMICNLETTYRSKKPATNLKMPGVYNVD<br>HRLRIKEADDETYVEQHEVAVARYSTGGAGDGGKSGSGSLEHHHHHHH |
| P3SN_4h-FusionRed | P3_4h-linker(32AA), labelling site-TEV site-linker(7AA)-FusionRed<br>(4-heptade P3SN coiled-coil (foothold) fused to FusionRed. Used for BLI.)<br>MGSPEDIQLEEEISQLEQKNSSELKEKNQELKYGGSGSGSGSGSGSGSGSGSGSG<br>SDEGCGEDGSENLYFQSGSGSGSGMVSELIKENMPMKLYMEGTVNNHHFKCTSEG<br>EGKPYEGTQTMRIKVVEGGPLPFAFDILATSFMYGSRTFIKHPPGIPDFFKQSFPEGF<br>TWERVTTYEDGGVLTATQDTSQDGLIYNVVKVRGVNFPANGPVMQKKTGWEA<br>STETMYPADGGLEGACDMALKLVGGGHMICNLETTYRSKKPATNLKMPGVYNVD<br>HRLRIKEADDETYVEQHEVAVARYSTGGAGDGGKSGSGSLEHHHHHHH |
| P4SN-FusionRed | P4-linker(32AA), labelling site-TEV site-linker(7AA)-FusionRed |
|  | (4-heptade P4SN coiled-coil (foot) fused to FusionRed. Used for BLI.)<br>MGSPEDKISQLKEKIQQLKQENQQLEENSQLEYGGSGGGSGGGSGGGSGSGSGSGS<br>GSDEGCGEDGSENLYFQGS GSGSGSSGMVSELIKENMPMKLYMEGTVNNHHFKCTSE<br>GEGKPYEGTQTMRIKVVEGGPLPFAFDILATSFMYGSRTFIKHPPGIPDFFKQSFPEG<br>FTWERTTYEDGGVLTATQDTSLQDGLIYNVKVRGVNFPANGPVMQKKTLGWE<br>ASTETMYPADGGLEGACDMALKLVGGGHMICNLETTYRSKKPATNLKMPGVYNV<br>DHRLERIKEADDETYVEQHEVAVARYSTGGAGDGGKGS GSGSLEHHHHHH |

**Extended Data Table 3:** Overview of the final walker designs.

| Walker | Full name | Stable at low concentrations | Oligomeric state | Feet distance [nm] | Feet attachment terminus | Reference |
| --- | --- | --- | --- | --- | --- | --- |
| WALKER-C4 | C4-71-8-1_P4mSN | yes | 4 | 5.8 | N | <sup>16</sup> |
| WALKER-C4b | C4-80-4-1_2R_P4mSN | no | 4 | 4.8 | C | <sup>16</sup> |
| WALKER-C5 | C5_HF_0019_P4mSN | yes | 5 | 5.5 | N | <sup>15</sup> |
| WALKER-C6 | C6_1BH_11_P4mSN | yes | 6 | 5.0 | N | <sup>15</sup> |
| WALKER-C6b | C6_80_6H2LD_P4mSN | yes | 6 | 5.6 | N | <sup>15</sup> |
| WALKER-C8 | C8-71-7-1_P4mSN | yes | 8 | 5.5 | N | <sup>16</sup> |

**Extended Data Table 4:** Diffusion rates and dwell times of walkers on both types of tracks with mNeonGreen fusion. Data is represented as mean ± s.d.

|  | TRACK-P3SN_3.5h-mNG |  |  | TRACK-P3SN_4h-mNG |  |  |
| --- | --- | --- | --- | --- | --- | --- |
| WALKER | D<sub>e1</sub> [nm<sup>2</sup>/s] | D<sub>e2</sub> [nm<sup>2</sup>/s] | Dwell time [s] | D<sub>e1</sub> [nm<sup>2</sup>/s] | D<sub>e2</sub> [nm<sup>2</sup>/s] | Dwell time [s] |
| WALKER-C4 | 1030 ± 330 | 510 ± 230 | 25 ± 17 | 1140 ± 440 | 420 ± 170 | 79 ± 76 |
| WALKER-C5 | 1190 ± 470 | 540 ± 170 | 59 ± 53 | 1470 ± 450 | 560 ± 200 | 57 ± 79 |
| WALKER-C6 | 1510 ± 390 | 480 ± 100 | 25 ± 29 | 1650 ± 500 | 670 ± 210 | 66 ± 84 |
| WALKER-C6b | 1380 ± 450 | 570 ± 180 | 28 ± 26 | 1790 ± 690 | 550 ± 230 | 73 ± 60 |
| WALKER-C8 | 1560 ± 380 | 640 ± 210 | 61 ± 89 | 1890 ± 720 | 550 ± 170 | 79 ± 148 |
| P4SN_4h | | | | 290 ± 60 | 210 ± 30 | 14 ± 5 |
| immobilized beads | | | | 22 ± 2 | 15 ± 2 | |

**Extended Data Table 5:** Average lengths of steps of walkers on both types of tracks with mNeonGreen fusion. Data is represented as mean ± s.d.

|  | TRACK-P3SN_3.5h-mNG | TRACK-P3SN_4h-mNG |
| --- | --- | --- |
| WALKER | Step length [nm] | Step length [nm] |
| WALKER-C4 | 29 ± 23 | 28 ± 22 |
| WALKER-C5 | 30 ± 21 | 32 ± 23 |
| WALKER-C6 | 30 ± 22 | 33 ± 23 |
| WALKER-C6b | 31 ± 23 | 34 ± 25 |
| WALKER-C8 | 33 ± 24 | 31 ± 23 |

**Extended Data Table 6:** Diffusion rates of walkers C4 and C8 on both types of tracks without mNeonGreen at different concentrations of sucrose and temperatures. Data is represented as mean ± s.d.

|  |  | Diffusion coefficient [nm<sup>2</sup>/s] |  |  |  |
| --- | --- | --- | --- | --- | --- |
|  |  | TRACK-P3SN_3.5h |  | TRACK-P3SN_4h |  |
| WALKER | CONDITIONS | D<sub>e1</sub> [nm<sup>2</sup>/s] | D<sub>e2</sub> [nm<sup>2</sup>/s] | D<sub>e1</sub> [nm<sup>2</sup>/s] | D<sub>e2</sub> [nm<sup>2</sup>/s] |
| WALKER-C4 | 25 °C, 0 % sucrose, 75 mM NaCl | 740 ± 180 | 390 ± 60 | 1010 ± 280 | 390 ± 60 |
| WALKER-C8 | 25 °C, 0 % sucrose, 75 mM NaCl | 1610 ± 410 | 640 ± 230 | 1840 ± 510 | 720 ± 170 |
| WALKER-C8 | 45 °C, 0 % sucrose, 75 mM NaCl | 2700 ± 850 | 1120 ± 121 | 2610 ± 630 | 1250 ± 230 |
| WALKER-C8 | 25 °C, 20 % sucrose, 75 mM NaCl | 1620 ± 450 | 630 ± 140 | 2080 ± 810 | 810 ± 310 |
| WALKER-C8 | 25 °C, 50 % sucrose, 75 mM NaCl | 820 ± 380 | 340 ± 120 | 530 ± 140 | 280 ± 50 |
| WALKER-C8 | 25 °C, 50 % sucrose, 500 mM NaCl | 960 ± 460 | 340 ± 120 | 1200 ± 510 | 390 ± 130 |

**Extended Data Table 7:**
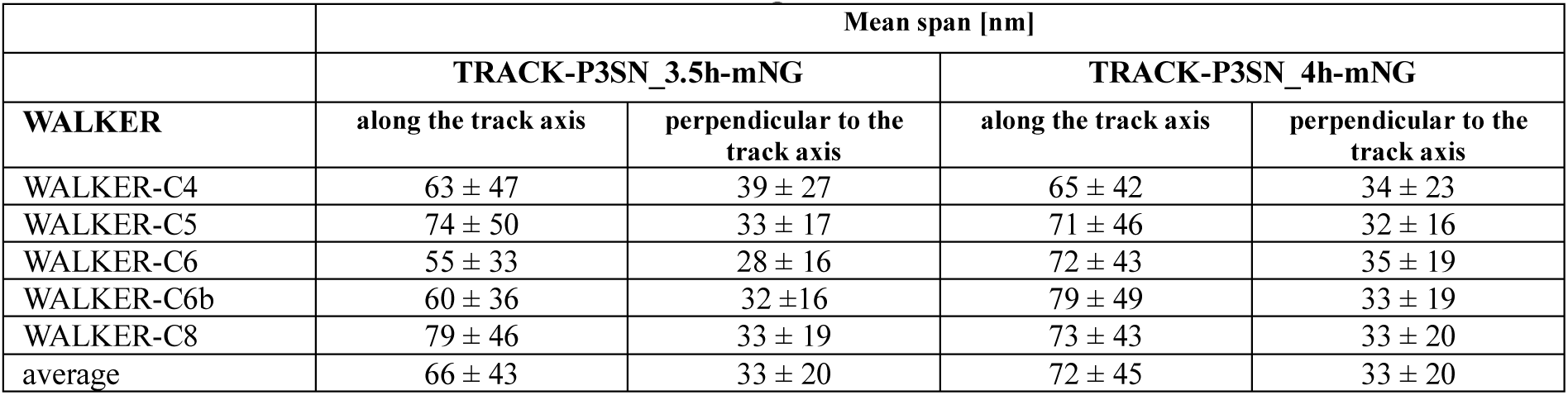
Average trajectory spans of walkers along and perpendicular to the track axis. Calculated for tracks with mNeonGreen. Data is represented as mean ± s.d.

|  | Mean span [nm] |  |  |  |
| --- | --- | --- | --- | --- |
|  | TRACK-P3SN_3.5h-mNG |  | TRACK-P3SN_4h-mNG |  |
| WALKER | along the track axis | perpendicular to the track axis | along the track axis | perpendicular to the track axis |
| WALKER-C4 | 63 ± 47 | 39 ± 27 | 65 ± 42 | 34 ± 23 |
| WALKER-C5 | 74 ± 50 | 33 ± 17 | 71 ± 46 | 32 ± 16 |
| WALKER-C6 | 55 ± 33 | 28 ± 16 | 72 ± 43 | 35 ± 19 |
| WALKER-C6b | 60 ± 36 | 32 ± 16 | 79 ± 49 | 33 ± 19 |
| WALKER-C8 | 79 ± 46 | 33 ± 19 | 73 ± 43 | 33 ± 20 |
| average | 66 ± 43 | 33 ± 20 | 72 ± 45 | 33 ± 20 |

**Extended Data Table 8:**
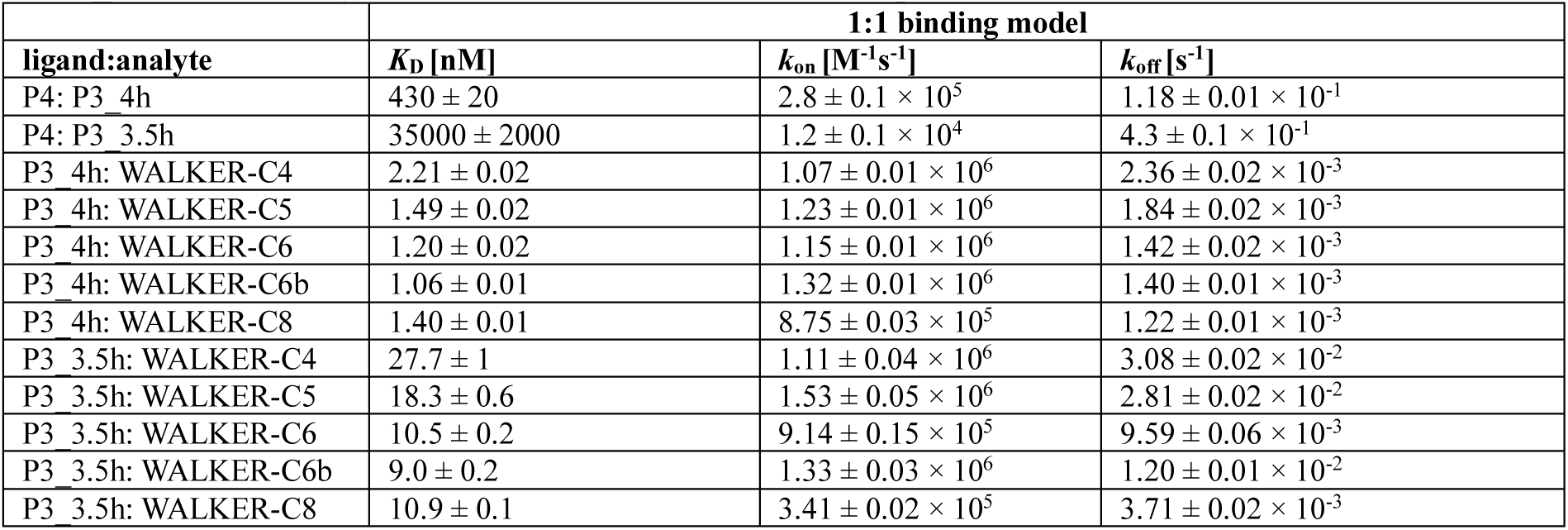
Kinetic parameters for walker and single foot binding to P3SN_3.5h and P3SN_4h, measured by BLI at sensor saturation. Data is represented as mean ± s.e.m.

|  | 1:1 binding model |  |  |
| --- | --- | --- | --- |
| ligand:analyte | $K_D$ [nM] | $k_{on}$ [M<sup>-1</sup>s<sup>-1</sup>] | $k_{off}$ [s<sup>-1</sup>] |
| P4: P3_4h | 430 ± 20 | 2.8 ± 0.1 $\times 10^5$ | 1.18 ± 0.01 $\times 10^{-1}$ |
| P4: P3_3.5h | 35000 ± 2000 | 1.2 ± 0.1 $\times 10^4$ | 4.3 ± 0.1 $\times 10^{-1}$ |
| P3_4h: WALKER-C4 | 2.21 ± 0.02 | 1.07 ± 0.01 $\times 10^6$ | 2.36 ± 0.02 $\times 10^{-3}$ |
| P3_4h: WALKER-C5 | 1.49 ± 0.02 | 1.23 ± 0.01 $\times 10^6$ | 1.84 ± 0.02 $\times 10^{-3}$ |
| P3_4h: WALKER-C6 | 1.20 ± 0.02 | 1.15 ± 0.01 $\times 10^6$ | 1.42 ± 0.02 $\times 10^{-3}$ |
| P3_4h: WALKER-C6b | 1.06 ± 0.01 | 1.32 ± 0.01 $\times 10^6$ | 1.40 ± 0.01 $\times 10^{-3}$ |
| P3_4h: WALKER-C8 | 1.40 ± 0.01 | 8.75 ± 0.03 $\times 10^5$ | 1.22 ± 0.01 $\times 10^{-3}$ |
| P3_3.5h: WALKER-C4 | 27.7 ± 1 | 1.11 ± 0.04 $\times 10^6$ | 3.08 ± 0.02 $\times 10^{-2}$ |
| P3_3.5h: WALKER-C5 | 18.3 ± 0.6 | 1.53 ± 0.05 $\times 10^6$ | 2.81 ± 0.02 $\times 10^{-2}$ |
| P3_3.5h: WALKER-C6 | 10.5 ± 0.2 | 9.14 ± 0.15 $\times 10^5$ | 9.59 ± 0.06 $\times 10^{-3}$ |
| P3_3.5h: WALKER-C6b | 9.0 ± 0.2 | 1.33 ± 0.03 $\times 10^6$ | 1.20 ± 0.01 $\times 10^{-2}$ |
| P3_3.5h: WALKER-C8 | 10.9 ± 0.1 | 3.41 ± 0.02 $\times 10^5$ | 3.71 ± 0.02 $\times 10^{-3}$ |

**Extended Data Table 9:** Kinetic parameters of WALKER-C8 binding to P3SN_3.5h and P3SN_4h at different loading concentrations. Data is represented as mean ± s.e.m.

|  |  | 1:1 binding model |  |  |
| --- | --- | --- | --- | --- |
| ligand:analyte | loading concentration [nM] | $K_D$ [nM] | $k_{on}$ [M<sup>-1</sup>s<sup>-1</sup>] | $k_{off}$ [s<sup>-1</sup>] |
| P3_4h: WALKER-C8 | 75 | 1.40 ± 0.01 | 8.75 ± 0.03 $\times 10^5$ | 1.22 ± 0.01 $\times 10^{-3}$ |
| P3_4h: WALKER-C8 | 10 | 1.87 ± 0.2 | 1.06 ± 0.00 $\times 10^6$ | 1.99 ± 0.02 $\times 10^{-3}$ |
| P3_4h: WALKER-C8 | 5 | 2.70 ± 0.03 | 1.22 ± 0.01 $\times 10^6$ | 3.30 ± 0.02 $\times 10^{-3}$ |
| P3_4h: WALKER-C8 | 1 | 22 ± 0.9 | 2.61 ± 0.10 $\times 10^6$ | 5.75 ± 0.07 $\times 10^{-2}$ |
| P3_3.5h: WALKER-C8 | 75 | 10.9 ± 0.1 | 3.41 ± 0.02 $\times 10^5$ | 3.71 ± 0.02 $\times 10^{-3}$ |
| P3_3.5h: WALKER-C8 | 10 | 41.5 ± 1.0 | 4.75 ± 0.11 $\times 10^5$ | 1.97 ± 0.01 $\times 10^{-2}$ |
| P3_3.5h: WALKER-C8 | 5 | 180 ± 10 | 6.79 ± 0.30 $\times 10^5$ | 1.22 ± 0.02 $\times 10^{-1}$ |
| P3_3.5h: WALKER-C8 | 1 | / | / | / |

## Methods

### Design of large diameter HB67 fibres

The helical docking and design strategy^27^ was applied to proteins (DHR04, DHR05, DHR07, DHR08, DHR14, DHR53, DHR54, DHR64, DHR71, DHR76, DHR79, DHR81, ZCON15, ZCON37, ZCON127, ZCON131, pRO-2.3) in order to generate models of helical filament assemblies. A wider range of cyclic and N jumps combination were sampled to get fibres with larger diameter. Candidate designs were filtered using several metrics: a Rosetta energy difference of less than –15.0 between the polymerized (bound) and monomeric (unbound) states, an interface area larger than 700 Å², a shape complementarity score above 0.62, and no more than five unsatisfied polar residues. Designs passing these benchmarks were subjected to manual refinement, during which non-critical mutations were reverted to their native sequence. For each configuration, the top-scoring model was chosen and carried forward into the final protein set for experimental validation.

### Design of H-fused fibres

The HB67 fibre was modified on the C-terminal using the Rosetta MergePDB mover. The following repeat proteins (DHRs^10^) were used as possible fusion blocks: CP15_4, DHR01, DHR03, DHR04, DHR05, DHR07, DHR08, DHR09, DHR09-xtal_4, DHR10, DHR14_hp_repack, DHR15, DHR18, DHR20, DHR21, DHR23, DHR24, DHR26, DHR27, DHR31, DHR32, DHR36, DHR39, DHR46, DHR47, DHR49_4repeat_XtalFit, DHR49, DHR52, DHR53, DHR54, DHR55, DHR57, DHR58, DHR59, DHR62, DHR64, DHR68, DHR70, DHR71, DHR72, DHR76, DHR77, DHR78, DHR79, DHR80, DHR81, DHR82, PEP12_4, PEP12_6, PX3_4, PXX2_4, PXX13_4, PXX28_4.

The following overlap lengths (and maximal allowed RMSD in overlap) were:

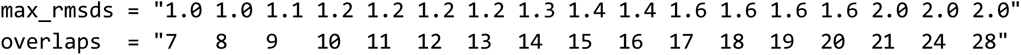

For details see 01_fuse_to_fibers_merge.xml in https://github.com/ajasja/fibre-H-fuse.

The side chains of the 452 passing fusion designs were redesigned using the Fast Design in symmetry mode. The interface of the HB67 fibre was fixed, to ensure that fibres will still from. During the design phase the protomers were expanded in symmetry mode and checked for clashes between symmetry units. For details see 02_design_fusions.xml.

Designs were further filtered using the following metrics:

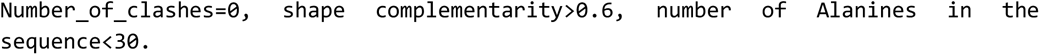

Designs were sorted by score_per_residue and following manual inspection in Pymol 2.5.0, 18 designs were ordered and named HA1 to HA18 (Extended Data Fig. 2). For details see 05_analyze_rescored.ipynb.

Rosetta version 2020.25+HEAD.d2d9f90b8cb^28^ was used for all calculations.

### Design of walkers

Homooligomers with fused DHR were selected from Hsia et al^15^ and Edman et al^16^ (Extended Data Table 3). The middle repeat unit was identified, and the number of middle repeat units was adjusted until the spacing between the repeats matched approximately 5 nm (matching the spacing on the fibre). The alignment of DHRs for extension and measurement was done manually in Pymol 2.5.0. Models with flexibly attached feet were constructed by predicting the sequence using ColabFold 1.4.0^29^.

### Expression and purification of HB67, HA4 and HA13 fibres

Synthetic genes were codon-optimized for *Escherichia coli* expression and synthesized by IDT. Constructs were inserted into pET29b+ vectors, incorporating N- or C-terminal GFP fusions where space permitted, using the NdeI and XhoI restriction sites. Plasmids were transformed into BL21* (DE3) *E. coli* cells, which were cultured in 50 mL Terrific Broth supplemented with kanamycin (200 mg L⁻¹). Protein expression was induced using the Studier autoinduction protocol^30^ at 37 °C for 24 h. Following induction, cells were collected by centrifugation, resuspended in Tris-buffered saline (TBS), and lysed with BugBuster reagent. The soluble fraction was clarified by centrifugation and purified by Ni²⁺-IMAC using Ni-NTA Superflow resin. Resin-bound proteins were washed with ten column volumes of buffer containing 40 mM imidazole and 500 mM NaCl, and eluted with buffer containing 400 mM imidazole and 75 mM NaCl. Soluble and insoluble fractions were analysed by SDS–PAGE, and constructs showing bands at the expected molecular weight were advanced for electron microscopy screening. For extended characterization, selected designs were scaled up to 0.5 L cultures, expressed under identical conditions, harvested after 24 h at 37 °C, and processed by microfluidization and purification following the same workflow.

### Negative stain electron microscopy screening

Soluble protein fractions were concentrated into buffer (25 mM Tris, 75 mM NaCl, pH 8) for negative-stain electron microscopy. For sample preparation, 6 µL drops were applied to carbon-coated 200-mesh copper grids that had been glow-discharged; each drop consisted of 1 µL sample diluted immediately with 5 µL buffer. Grids were washed with Milli-Q water and stained with 0.75% uranyl formate (pH 4.0), following established procedures^31^. Screening was carried out on a Talos L120C transmission electron microscope (Thermo Fisher) operating at 120 kV. Micrographs were recorded using a bottom-mounted Teitz CMOS 4k camera, and image contrast was enhanced using Fiji software^32^.

### Testing of reversible heterodimers on fibres

A backbone vector (pet29b_HA13LL_mNeonGreen.gb and pet29b _HA4LL_mNeonGreen.gb) was ordered at GenScript Biotech (NJ, USA). The heterodimers were ordered as synthetic genes and cloned into the backbone vector by Genscript using their VectorArk Cloning technology. The constructs were assembled form the following parts: HB67-DHR-linker(10AA)-**HETERODIMER**-linker(34AA)-labelling site-TEV site-mNG-linker(6AA)-His6.

Tested heterodimers are described in Extended Data Table 1.

The constructs were expressed and purified as described in “Expression and purification of HB67 and HA13 fibres”. Fibres were dialysed overnight in 3ml 3.5 kDa dialysis cassettes. Next samples were diluted (10x, 100x and 1000x) and screened in a Greiner Bio-one SensoPlate 96 well glass bottom plate (Greiner Bio-one, catalog #655892) using an IN Cell Analyzer 2500HS (Molecular Devices) and a 60× 0.95 NA CFI Plan Apo objective (Nikon). Cells were illuminated with a seven-color Solid State Illuminator (SSI) for fluorescence excitation, specifically a 473 nm excitation LED at 20 ms exposure time. Fluorescence signals were acquired sequentially using the following filters: Green (excitation 473/28 nm, emission 511/23 nm), yellow/orange (excitation nm, emission nm), and far-red (excitation 631/28 nm, emission 684/24 nm). Imaging was controlled using the IN Cell Analyzer 2500 HS software version 7.4 and light collected on sCMOS camera without binning. Samples were evaluated visually for fibre formation.

### Track and walker protein expression and purification

All walker and tracks with mNeonGreen constructs were ordered from GenScript Biotech (NJ, USA) already inserted into pET29b+ expression vector. Gibson assembly was used to delete the mNeonGreen fusion from the track constructs. The P4SN and P3SN constructs fused to FusionRed – used for kinetic characterization – were also ordered from GenScript, inserted into pET29b+ vector. Gibson assembly was used to delete three aminoacids (E7, I8, Q9) in P3SN to create the 3.5hP3SN-FusionRed construct.

The plasmids were transformed into NiCO21 (DE3) production strain (NEB, MA, USA) and grown overnight at 37 °C on LB agar plates containing kanamycin (50 μg/mL). Single colonies were picked from the plates and inoculated into 10 mL LB medium supplemented with kanamycin (50 μg/mL). The cultures were grown overnight at 37 °C. 4 mL of the overnight culture were inoculated into 200 mL of LB medium supplemented with kanamycin (50 μg/mL) and then incubated at 37 °C. When OD_600_ of the culture reached ∼0.7, IPTG (Goldbio, MO, USA) was added to 0.4 mM final concentration. Cultures were incubated overnight at 20 °C, after which the cells were harvested by centrifugation. The pellets were resuspended in Tris buffer (20 mM Tris pH 7.5, 75 mM NaCl for tracks and 150 mM NaCl for walkers and feet, 1 mM TCEP) supplemented with 20 mM imidazole, protease inhibitor cocktail (CPI, Millex Sigma-Aldrich, MO, USA) and 15 U/mL Benzonase (Merck, Germany), followed by lysis by sonication. The lysates were centrifuged for 30 min at 15000 rcf and 4 °C. The supernatants of walker proteins and feet were filtered through a 0.22 µm filter (Minisart, Sartorius, Germany), whereas track supernatants were not so as to prevent the loss of protein. All proteins were isolated and purified on gravity columns with 1 mL of Ni-NTA resin, pre-equilibrated with Tris buffer (20 mM Tris pH 7.5, 75 mM NaCl for tracks and 150 mM NaCl for walkers and feet, 1 mM TCEP) supplemented with 20 mM imidazole. The washing steps were carried out with equilibration buffer supplemented with 20 mM imidazole, followed by washing with equilibration buffer supplemented with 30 mM imidazole. Proteins were eluted from the resin with equilibration buffer supplemented with 250 mM imidazole.

Directly after elution glycerol was added to tracks to the final concentration of 5 % (w/v), followed by plunge freezing in liquid nitrogen. The tracks were stored at −80 °C at ∼ 15 μM concentration. The feet fused to FusionRed were dialysed overnight against Tris buffer (20 mM Tris pH 7.5, 150 mM NaCl, 1 mM TCEP) and stored at −80 °C at ∼ 250 μM concentration.

Walkers were additionally purified by size-exclusion chromatography, using a Superdex 200 Increase 10/300 GL column (Cytiva, MA, USA). The mobile phase was Tris buffer (20 mM Tris pH 7.5, 150 mM NaCl, 1 mM TCEP) supplemented with 5 % (w/v) glycerol. Fractions containing protein were merged and concentrated to ∼ 1 mg/mL, followed by plunge freezing in liquid nitrogen and storing at −80 °C.

### Size Exclusion Chromatography Coupled to Multi-angle Light Scattering

150 μL aliquots of walker proteins were thawed and filtered through 0.1 μm centrifuge filters (Merck Millipore, MA, USA) and subsequently injected onto a Superdex 200 Increase 10/300 GL column (Cytiva, MA, USA), coupled to a Waters e2695 high-performance liquid chromatography system, equipped with a UV, refractive index and multi-angle light scattering detectors. BSA with concentration 1 mg/mL was used as the internal standard at the start and end of the sequence of sample runs. The mobile phase was Tris buffer (20 mM Tris pH 7.5, 150 mM NaCl, 1 mM TCEP). Results were analysed with Astra 7.0 software (Wyatt, CA, USA).

### Mass photometry (MP)

Mass photometry measurements on walker proteins were carried out with a Refeyn Two^MP^ mass photometer (Oxford, UK). Walkers were diluted in Tris buffer (20 mM Tris pH 7.5, 150 mM NaCl, 1 mM TCEP) to the lowest concentration that still produced enough counts (1000 counts minimum) during the measurement, ranging from 20 – 40 nM, depending on the walker. The walkers were measured either directly after dilution, 2 hours after dilution or after incubation at 95 °C for 2 hours. Standard mass photometry uncoated sample carrier glasses and casettes (Refeyn, Oxford, UK) were used for the measurement. Each sample was measured for 60 seconds. For determination of mass MassFerence P1 Calibrant (Refeyn, Oxford, UK) was used as an internal standard. The measurements were analysed with Discover^MP^ software.

### Isothermal calorimetry (ITC)

Peptides were synthesized by Proteogenix (France) and solubilized in a modified SEC buffer (20 mM TRIS-HCl, 500 mM NaCl, and 1 mM TCEP). Peptide concentrations were determined by measuring absorbance at 280 nm. Isothermal titration calorimetry (ITC) was conducted using MicroCal VP-ITC (Malvern Instruments, UK), with 28 injections of 10 µL each titrated into a 2 mL sample cell. Peptide P4SN_4h (at concentrations of 3 µM and 15 µM) was in the cell as the analyte, while P3SN_4h (30 µM) and P3SN_3.5h (150 µM) served as the titrants. A 600-second equilibration period followed each injection. Peak integration was performed using the NITPIC software (ref) and analysis was done SEDPHAT.

### Biolayer interferometry (BLI)

For BLI measurements, the feet and footholds (P4SN, P3SN_4h and P3SN_3.5h) were flexibly fused to FusionRed protein to enhance their mass and consequently the signal measured with BLI. P4SN was used as the ligand for single feet measurements, while P3SN_3.5h and P3SN_4h were used for walker measurements. To attach the ligands to the sensor, the proteins were biotinylated with EZ-Link maleimide-PEG2-biotin (Thermo Fischer Scientific, CA, USA) through a reduced cysteine residue. The proteins were mixed with maleimide-PEG2-biotin in a 1:20 ratio and incubated overnight at 4 °C. The unreacted maleimide-PEG2-biotin was removed with the use of 0.5 mL Zeba spin desalting column with 7 kDa MWCO (Thermo Fischer Scientific, CA, USA).

The BLI measurements were performed with Octet R4 Protein Analysis System (Sartorius, Germany) and High Precision Streptavidin (SAX) sensors were used. All samples were prepared in Tris buffer (20 mM Tris pH 7.5, 150 mM NaCl, 1 mM TCEP). To block nonspecific binding to the sensors, 0.5 % (w/v) BSA was included in all steps of the assay except for the first baseline measurement. In single feet experiments, ligand scout was performed beforehand to determine the appropriate concentration of ligand for the kinetic assay. A concentration of 10 nM P4SN was selected for measurements with P3SN_4h, and 50 nM for P3SN_3.5h. For measurements with P3SN_4h, two assays were performed with the following analyte concentrations: 5450, 220, 870, 0 nM and 349, 140, 55.8 and 0 nM. The samples with 0 nM concentration of analyte served as references. For measurements with P3SN_3.5h, two assays were performed with the following analyte concentrations: 200, 150, 100, 0 µM and 75, 50, 25 and 0 µM. Due to the high analyte concentrations and the resulting significant nonspecific binding, two additional assays were conducted using unloaded sensors at the same analyte concentrations. These served as reference sensors. The following step times were used for each measurement: 60 s for both baselines, 600 s for ligand loading, 45 s (P3SN_4h) and 60 s (P3SN_3.5h) for association, 60 s (P3SN_4h) and 45 s (P3SN_3.5h) for dissociation. Plate shaking was maintained at 1000 rpm throughout.

Walker measurements began with ligand-saturated sensors loaded with 75 nM of either P3SN_3.5h or P3SN_4h. For P3SN_3.5h, two assays were performed using walker protomer concentrations of 10, 5, 2, 0 µM and 1, 0.5, 0.25, 0 µM; for P3SN_4h, the protomer concentrations were 1, 0.5, 0.25, 0 µM and 0.125, 0.0625, 0.0313, 0 µM. Samples without analyte (0 µM) served as references. Except for association and dissociation steps (set to 100 s in all walker assays), timing matched the single foot protocol. For experiments using lower ligand loading concentrations, the same walker concentrations were applied. Ligand loading was performed with 10, 5, and 1 nM P3SN_4h, and 10 and 5 nM P3SN_3.5h. Step timings were identical to those in the saturated-sensor assays, except for loading, which was decreased to 120 s to prevent saturating the sensor, and the second baseline, which was prolonged to 250 s to ensure sufficient blocking of the sensor with BSA. Shaking speed remained constant at 1000 rpm.

The measurements for both pairs were analysed using the Octet Analysis software. First, the values of the reference samples were deducted from the measurements. For the single feet measurements with P3SN_3.5h ligand, the values of the reference sensors were also deducted. The Y-values of the measurements were then aligned to the average of the baseline step and the steps were aligned to the dissociation step. For the kinetic analysis, 1:1 binding model and global fitting were used. When analysing the walker measurements, oligomer concentrations were used instead of protomer concentrations. Based on SEC-MALS results (Fig. 3b), which confirmed a relatively homogeneous oligomer population, oligomer concentrations were estimated by dividing the total protomer concentration by the number of protomers per oligomer.

### Negative stain electron microscopy of tracks

Samples of fibres and tracks at a concentration of ∼ 1 mg/mL were vortexed, and 7 μL of the sample was pipetted onto a formvar/carbon grid (Ted Pella Inc, CA, USA, cat. 01753F). After a 5-minute incubation, the excess sample was blotted off with filter paper. The grid was then washed with a drop of ultrapure water, which was also blotted off. A drop of 1 % uranyl acetate was applied and immediately blotted off with filter paper. Once dry, the grids were stored at room temperature until imaging.

The samples were imaged with Apreo 2 Scanning Electron Microscope (Thermo Fischer Scientific, CA, USA) at 30 kV and 13 pA with a STEM 3+ detector in brightfield mode.

### Fibre Cryo-EM grid preparation and data collection

HA13 cryoEM samples were prepared by applying protein to CFLAT 2/2 holey-carbon grids (Protochips), blotting away liquid and plunging into liquid ethane using a Vitrobot (Thermofisher). Data was collected on a Titan Krios (Thermofisher) equipped with a Quantum Gatan Imaging Filter energy filter (Gatan Inc.) operating in zero-loss mode with a 20-eV slit width. Movies were acquired using a K-2 Summit Direct Detect camera (Gatan Inc.) operating in superresolution mode with a pixel size of 0.525 Å/pixel, with 50 frames and a total dose of 90 electrons/ Å^2^. Data collection was automated using Leginon^33^. The workflow for HA13 data processing is summarized in Extended Data Fig. 3. Movies were aligned, dose-weighted, and summed using the Relion^34^ implementation of MotionCor2^35^, with resulting micrographs binned to a pixel size of 1.05 Å/pixel. CTF parameters were estimated using Gctf^36^, and subsequent data processing was performed using Relion. Helices were picked manually from a subset of roughly 100 micrographs, then subjected to 2D classification to generate templates for subsequent automatic picking of helices from all micrographs. Particles from automatic helical picking were subjected to 2D classification, and high quality 2D class averages were selected for 3D refinement. A series of initial 3D helical refinements were performed with 4X binned particles (4.2 Å/pixel) and a cylinder starting model, using various helical symmetry values estimated from inspection of 2D class averages and their power spectra. An initial 3D refinement performed with a helical rise of 9Å and a helical rotation of 37° produced a structure with clear secondary structure. Inspection of this structure revealed additional C2 symmetry. This initial structure was used as a starting map for subsequent 3D helical refinement using unbinned particles, where helical rise and rotation were refined and C2 symmetry was imposed. Beamtilt, anisotropic magnification, and per-particle defocus were refined prior to a final 3D helical refinement. Density modification of the final structure was performed using ResolveCryoEM in Phenix^37,38^.

### Walker Cryo-EM data processing

For grid preparation, 3.5 µL of sample with concentrations between 0.5 and 1 mg/mL was applied to glow-discharged 300 mesh copper Quantifoil holey carbon grids (1.2/1.3), coated with graphene oxide. Grids were plunge-frozen using a Vitrobot Mark IV. Samples were then screened and imaged on a 200 kV Glacios transmission electron microscope equipped with a Falcon 3 camera. Movies were acquired at a nominal magnification of 190,000x with a total dose of approximately 40 e^−^ Å^−2^.

Data processing was performed in cryoSPARC. The general workflow included motion correction and CTF estimation, followed by particle picking and iterative 2D classification to remove junk particles. This was followed by ab initio reconstruction, and subsequent refinement. Ab initio reconstruction was performed with C1 symmetry, while refinements were carried out both with and without symmetry enforced. Imposing symmetry generally resulted in better-quality maps. Although all samples exhibited preferred orientation and had overall lower nominal resolution, we were still able to refine the structures to sufficient resolutions to confirm the general designs.

### Fluorescent labelling of walkers

All walker proteins contain one cysteine residue on their C-terminus, which was utilized for attaching a fluorescent label via a thiol-maleimide reaction. Walker and ATTO 643 maleimide (ATTO-TEC GmbH, Germany, cat. AD 643-41) were mixed in 1:3 molar ratio in Tris buffer (20 mM Tris pH 7.5, 150 mM NaCl, 1 mM TCEP) and incubated overnight at 4 °C, shielded from light. The protein was separated from unreacted dye with a PD-10 desalting column packed with Sephadex G-25 resin (Cytiva, MA, USA). Tris buffer (20 mM Tris pH 7.5, 150 mM NaCl, 1 mM TCEP) supplemented with 5 % glycerol was used a the mobile phase. The fractions containing both protein and dye were merged and concentrated to ∼ 30 μM. Aliquots were plunge frozen in liquid nitrogen and stored at −80 °C.

### Preparation of oxygen scavenger solutions

A mixture of glucose oxidase from *Aspergillus niger* (Sigma-Aldrich, MO, USA, cat. G7141) and catalase from bovine liver (Sigma-Aldrich, MO, USA, cat. C40) and Trolox (Sigma-Aldrich, MO, USA, cat. AL-238813-5G) were used as oxygen scavengers. 50x stock solution of enzyme mixture was prepared by mixing equal volumes of catalase and glucose oxidase solutions in Tris buffer (20 mM Tris pH 7.5, 75 mM NaCl, 1 mM TCEP) with concentrations 217000 U/mL and 16500 U/mL, respectively. The mixture was filtered through a 0.22 µm filter, aliquoted and stored at −80 °C until use.

100x stock of Trolox was prepared by adding 100 mg of Trolox to the following mixture: 430 μl 100 % Methanol, 69 μl 5 M NaOH and 3.47 ml dH_2_O. The suspension was aged overnight at room temperature and then filtered through a 0.22 µm filter. The stock was aliquoted and stored at −20 °C until use.

### Preparation of track sample for Total Internal Reflection Fluorescence Microscopy

The slides for TIRF microscopy were prepared as described previously^39^. Six channel Ibidi 1.5H glass coverslip slides (cat. 80607) were cleaned using 2 % Hellmanex (Sigma-Aldrich, MO, USA, cat. Z805939-1EA) solution and 10 M KOH. Next, 50 μL of PLL-*g*-PEG (SuSoS AG, Switzerland, cat. NB02-43) at a concentration of 0.3 mg/mL was flushed into the chamber, incubated for 30 min, and then washed with Tris buffer (20 mM Tris pH 7.5, 75 mM NaCl, 1 mM TCEP). Tracks were diluted to a concentration of ∼ 2 μM in Tris buffer (20 mM Tris pH 7.5, 75 mM NaCl, 1 mM TCEP) containing 0.5 % methylcellulose (viscosity 4000 cp, Sigma-Aldrich, MO, USA, cat. M0512) and deposited on the slides for 1 h, followed by washing 5 times with 200 μL Tris bufer (20 mM Tris pH 7.5, 75 mM NaCl, 1 mM TCEP).

### Preparation of negative control for single molecule tracking

For negative control, TetraSpeck beads (TetraSpeck Fluorescent Microspheres Sampler Kit, Thermo Fischer Scientific, CA, USA) with a diameter of 0.1 μm were diluted 1000x in ultrapure water and flown into a channel of a previously cleaned 6-channel Ibidi slide. The beads were left to dry overnight. Next, the slide was blocked and the tracks were deposited as described above.

### Single molecule tracking with Total Internal Reflection Fluorescence Microscopy

Walkers labelled with ATTO 643 were diluted to 0.25 nM in Tris buffer (20 mM Tris pH 7.5, 75 mM NaCl, 1 mM TCEP) supplemented with 1x oxygen scavenger mixture, 0.65 % glucose and 1x Trolox. When measuring diffusion at increased ionic strength, the concentration of NaCl in the buffer was increased to 500 mM. In case of single molecule tracking at increased viscosity, the buffer was also supplemented with 20 % (w/w) and 50 % (w/w) sucrose. Walkers were flowed into Ibidi slide channels containing tracks. The single molecule tracking experiments were performed on a LUMICKS C-Trap EDGE 350 microscope. The measurements were done immediately after the addition of walkers. For the negative control, measurements were performed with individual feet (P4SN_4h) diluted to 1 nM and TetraSpeck beads colocalized with the fibres. Both negative controls were performed with TRACK-P3SN_4h-mNG.

Tracks were imaged with IRM at 4 ms camera integration time and 40 fps with the averaging over 10 frames. Tracks containing mNeonGreen were also imaged with TIRFM. mNeonGreen was excited with 488 nm at 1 % laser power (0.015 mW) and a 525/40 emission filter was used. Walkers, feet and TetraSpecks were excited with 638 nm at 10% laser power (0.73 mW), and a 680/420 emission filter was used. In case of tracking at 45 °C, the laser power was increased to 30 % (2.2 mW) to compensate for loss of intensity due to spherical aberrations. Camera integration time was 200 ms and framerate was 4 fps.

### Preprocessing – color channel alignment and drift correction

Video output of the microscope consists of unaligned red, green, blue and IRM channels. To produce aligned videos, RGB channels were aligned through the Lumicks python package. The resulting TIFF stack consists of red, green, blue and the IRM channel.

To align the IRM channel it was necessary to calculate an alignment matrix. Pictures of 0.5 µm TetraSpeck beads (TetraSpeck Fluorescent Microspheres Sampler Kit, Thermo Fischer Scientific, CA, US) were taken in WT and IRM and the Picasso (version 0.6.8) Localize program was used to obtain the positions of the beads in the frame. Default parameters for the search were: LQ fit method, box size 21, minimum gradient 70000, maximum position error 3.5, but these parameters were adjusted as needed to optimize detection in different calibration images. With the positions of beads in both WT and IRM an affine transform matrix was calculated using the cv2 python package. The affine transform was applied to the IRM channel and all channels were merged into one image stack.

Drift correction of videos was performed using the Fast4DReg Fiji macro (https://github.com/CellMigrationLab/Fast4DReg). Default parameters were: time averaging 10, maximum expected drift 10, using the video first frame as reference. Due to significant mNeonGreen bleaching, the IRM channel was used as the reference for drift correction, both for tracks with mNeonGreen fusion and those without the fluorescent protein. Scripts are available at https://github.com/ajasja/LUMICKS_image_alignment.

### Diffusion analysis

To calculate the diffusion coefficients of the walkers the pre-processed TIFF stacks were processed further with the walker_tracker Python (version 3.10.6) pipeline (https://github.com/ajasja/walker_tracker) developed in this paper. First, a mask is created for each frame in the IRM channel using the scikit-image package (version 0.19.3), converting pixel values for tracks to 1 and all other pixel values to 0. This mask is then applied to all frames of the walker (red fluorescence) channel. The resulting red channel TIFF stack contains only walkers localized on tracks. The walkers in this stack are then localized using Picasso (version 0.6.8) Localize method with the following parameters: LQ fit method, box side length 7, minimum gradient 800, pixelsize 72 nm, drift 0. The resulting HDF5 file is opened as a dataframe using pandas (version 2.1.1) and further processed with the trackpy.link() function (trackpy package version 0.6.2) with the search range parameter set to 2 to identify and connect positions of walkers across frames to form trajectories. The length of each particle trajectory is calculated and trajectories shorter than 3 frames are filtered out. Sizes of steps in x and y direction are then calculated, and a two-dimensional histogram (numpy package version 1.24.3) of step sizes is generated, to which a two-dimensional Gaussian distribution is fit using curve_fit (scipy package version 1.11.2) and the following equation:

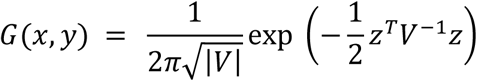

where 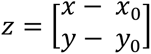 is the difference between the particle coordinates (x, y) and the mean (x_0_ , y_0_) and 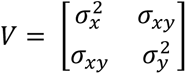 is the covariance matrix.

Variances along the principal axes – e_1_ and e_2,_ where e_1_ is the larger and e_2_ the smaller variance – are then calculated out of the resulting covariance matrix. The diffusion coefficients in each dimension are calculated with the following equation:

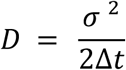

where σ^2^ represents e_1_ and e_2_.

For anisotropy analysis, TIFF stacks were manually cropped to include only straight track segments, then rotated to align the track axis with the y-axis. Walker trajectories within these stacks were recalculated using the same filtering parameters described above, and trajectory spans were determined.

For mean square displacement (MSD) analysis, trajectories from y-axis–aligned tracks were used. MSD was calculated with the trackpy.emsd() function, with max_lagtime set to 50 frames. To reduce noise, only trajectories longer than 50 frames were included.

## References

1. Yildiz, A. & Zhao, Y. Dyneins. Curr. Biol. 33, R1274–R1279 (2023).

2. Yildiz, A. Mechanism and regulation of kinesin motors. Nat. Rev. Mol. Cell Biol. 26, 86–103 (2025).

3. Trivedi, D. V., Nag, S., Spudich, A., Ruppel, K. M. & Spudich, J. A. The Myosin Family of Mechanoenzymes: From Mechanisms to Therapeutic Approaches. Annu. Rev. Biochem. 89, 667–693 (2020).

4. Abraham, Z., Hawley, E., Hayosh, D., Webster-Wood, V. A. & Akkus, O. Kinesin and Dynein Mechanics: Measurement Methods and Research Applications. J. Biomech. Eng. 140, (2018).

5. Watson, J. L. et al. De novo design of protein structure and function with RFdiffusion. Nature 620, 1089–1100 (2023).

6. Nilsson, P. et al. Tumbleweed: an artificial motor protein that walks along a DNA track. 2025.03.07.641123 Preprint at 10.1101/2025.03.07.641123 (2025).

7. Courbet, A. et al. Computational design of mechanically coupled axle-rotor protein assemblies. 9 (2022).

8. Shen, H. et al. De novo design of self-assembling helical protein filaments. Science 362, 705–709 (2018).

9. Hsia, Y. et al. Design of multi-scale protein complexes by hierarchical building block fusion. Nat. Commun. 12, 2294 (2021).

10. Brunette, T. J. et al. Exploring the repeat protein universe through computational protein design. Nature 528, 580–584 (2015).

11. Baryshev, A. et al. Massively parallel measurement of protein–protein interactions by sequencing using MP3-seq. Nat. Chem. Biol. 20, 1514–1523 (2024).

12. Chen, Z. et al. Programmable design of orthogonal protein heterodimers. Nature 565, 106–111 (2019).

13. Gradišar, H. & Jerala, R. *De novo* design of orthogonal peptide pairs forming parallel coiled-coil heterodimers. J. Pept. Sci. 17, 100–106 (2011).

14. Sahtoe, D. D. et al. Reconfigurable asymmetric protein assemblies through implicit negative design. Science 375, eabj7662 (2022).

15. Hsia, Y. et al. Design of multi-scale protein complexes by hierarchical building block fusion. Nat. Commun. 12, 2294 (2021).

16. Edman, N. I. et al. Modulation of FGF pathway signaling and vascular differentiation using designed oligomeric assemblies. Cell 187, 3726–3740.e43 (2024).

17. Schnitzbauer, J., Strauss, M. T., Schlichthaerle, T., Schueder, F. & Jungmann, R. Super-resolution microscopy with DNA-PAINT. Nat. Protoc. 12, 1198–1228 (2017).

18. Allan, D. B., Caswell, T., Keim, N. C., van der Wel, C. M. & Verweij, R. W. soft-matter/trackpy: v0.7. Zenodo 10.5281/zenodo.16089574 (2025).

19. Translational Diffusion of Globular Proteins in the Cytoplasm of Cultured Muscle Cells. Biophys. J. 78, 901–907 (2000).

20. Lu, H., Ali, M. Y., Bookwalter, C. S., Warshaw, D. M. & Trybus, K. M. Diffusive Movement of Processive Kinesin-1 on Microtubules. Traffic Cph. Den. 10, 1429–1438 (2009).

21. Shammas, S. L., Crabtree, M. D., Dahal, L., Wicky, B. I. M. & Clarke, J. Insights into Coupled Folding and Binding Mechanisms from Kinetic Studies *. J. Biol. Chem. 291, 6689–6695 (2016).

22. Bartoš, L., Lund, M. & Vácha, R. Enhanced diffusion through multivalency. Soft Matter 21, 179–185 (2025).

23. Weiss, M., Elsner, M., Kartberg, F. & Nilsson, T. Anomalous Subdiffusion Is a Measure for Cytoplasmic Crowding in Living Cells. Biophys. J. 87, 3518–3524 (2004).

24. Platani, M., Goldberg, I., Lamond, A. I. & Swedlow, J. R. Cajal body dynamics and association with chromatin are ATP-dependent. Nat. Cell Biol. 4, 502–508 (2002).

25. Smith, P. R., Morrison, I. E., Wilson, K. M., Fernández, N. & Cherry, R. J. Anomalous diffusion of major histocompatibility complex class I molecules on HeLa cells determined by single particle tracking. Biophys. J. 76, 3331–3344 (1999).

26. Weeramange, C. J., Fairlamb, M. S., Singh, D., Fenton, A. W. & Swint-Kruse, L. The strengths and limitations of using biolayer interferometry to monitor equilibrium titrations of biomolecules. Protein Sci. 29, 1004–1020 (2020).

27. Shen, H. et al. De novo design of self-assembling helical protein filaments. Science 362, 705–709 (2018).

28. Leaver-Fay, A. et al. ROSETTA3: an object-oriented software suite for the simulation and design of macromolecules. Methods Enzymol. 487, 545–574 (2011).

29. Mirdita, M. et al. ColabFold: making protein folding accessible to all. Nat. Methods 19, 679–682 (2022).

30. Studier, F. W. Protein production by auto-induction in high density shaking cultures. Protein Expr. Purif. 41, 207–234 (2005).

31. Nannenga, B. L., Iadanza, M. G., Vollmar, B. S. & Gonen, T. Overview of electron crystallography of membrane proteins: crystallization and screening strategies using negative stain electron microscopy. Curr. Protoc. Protein Sci. Chapter 17, Unit17.15 (2013).

32. Schindelin, J., et al. Fiji: an open-source platform for biological-image analysis. Nat. Methods 9, 676–682 (2012).

33. Suloway, C. et al. Automated molecular microscopy: the new Leginon system. J. Struct. Biol. 151, 41–60 (2005).

34. Scheres, S. H. W. RELION: implementation of a Bayesian approach to cryo-EM structure determination. J. Struct. Biol. 180, 519–530 (2012).

35. Zheng, S. Q. et al. MotionCor2: anisotropic correction of beam-induced motion for improved cryo-electron microscopy. Nat. Methods 14, 331–332 (2017).

36. Zhang, K. Gctf: Real-time CTF determination and correction. J. Struct. Biol. 193, 1–12 (2016).

37. Terwilliger, T. C., Ludtke, S. J., Read, R. J., Adams, P. D. & Afonine, P. V. Improvement of cryo-EM maps by density modification. Nat. Methods 17, 923–927 (2020).

38. Adams, P. D. et al. PHENIX: a comprehensive Python-based system for macromolecular structure solution. Acta Crystallogr. D Biol. Crystallogr. 66, 213–221 (2010).

39. Mezgec, K. et al. Coupling of Spectrin Repeat Modules for the Assembly of Nanorods and Presentation of Protein Domains. ACS Nano 18, 28748–28763 (2024).

